# An *Escherichia coli* ST131 pangenome atlas reveals population structure and evolution across 4,071 isolates

**DOI:** 10.1101/719583

**Authors:** Arun Gonzales Decano, Tim Downing

**Author notes:** School of Medicine, University of St. Andrews, UK.

## Abstract

*Escherichia coli* ST131 is a major cause of infection with extensive antimicrobial resistance (AMR) facilitated by widespread beta-lactam antibiotic use. This drug pressure has driven extended-spectrum beta-lactamase (ESBL) gene acquisition and evolution in pathogens, so a clearer resolution of ST131’s origin, adaptation and spread is essential. *E. coli* ST131’s ESBL genes are typically embedded in mobile genetic elements (MGEs) that aid transfer to new plasmid or chromosomal locations, which are mobilised further by plasmid conjugation and recombination, resulting in a flexible ESBL, MGE and plasmid composition with a conserved core genome. We used population genomics to trace the evolution of AMR in ST131 more precisely by extracting all available high-quality Illumina HiSeq read libraries to investigate 4,071 globally-sourced genomes, the largest ST131 collection examined so far. We applied rigorous quality-control, genome *de novo* assembly and ESBL gene screening to resolve ST131’s population structure across three genetically distinct Clades (A, B, C) and abundant subclades from the dominant Clade C. We reconstructed their evolutionary relationships across the core and accessory genomes using published reference genomes, long read assemblies and k-mer-based methods to contextualise pangenome diversity. The three main C subclades have co-circulated globally at relatively stable frequencies over time, suggesting attaining an equilibrium after their origin and initial rapid spread. This contrasted with their ESBL genes, which had stronger patterns across time, geography and subclade, and were located at distinct locations across the chromosomes and plasmids between isolates. Within the three C subclades, the core and accessory genome diversity levels were not correlated due to plasmid and MGE activity, unlike patterns between the three main clades, A, B and C. This population genomic study highlights the dynamic nature of the accessory genomes in ST131, suggesting that surveillance should anticipate genetically variable outbreaks with broader antibiotic resistance levels. Our findings emphasise the potential of evolutionary pangenomics to improve our understanding of AMR gene transfer, adaptation and transmission to discover accessory genome changes linked to novel subtypes.

## Background

Infections caused by multidrug-resistant (MDR) *Escherichia coli* sequence type (ST) 131 are increasing worldwide [1,2]. ST131 are extraintestinal pathogenic *E. coli* (ExPEC) associated with bloodstream and urinary tract infections and typically possessing extended-spectrum beta-lactamase (ESBL) genes [3–5], or more rarely carbapenemase genes [6]. MDR ST131 is a major cause of ExPEC infection because it has an extensive range of virulence factors [7–11] and may be highly pathogenic to hosts. ST131 has been reported in healthcare and community settings around the globe, and its dominant lineage Clade C is fluoroquinolone-resistant (FQ-R) [12,13]. Clade C has a type 1 fimbrial adhesin gene *H30* variant (*fimH*30) [10,14] and can offset the fitness costs of antimicrobial resistance (AMR), plasmid acquisition and maintenance through compensatory mutations at regulatory regions in contrast to FQ-susceptible Clades A and B [15].

Historically, *E. coli* population structure was inferred from allelic variation at seven housekeeping genes to assign ST complexes via multi-locus sequence typing (MLST) [16], or at 51 ribosomal genes for rST (ribosomal MLST) [17]. Outbreak investigation necessitates sufficient biomarker density to allow isolate discrimination, which is only possible with genome sequencing to allow profiling of all AMR genes [18,19]. Recent work applied core genome MLST (cgMLST) of 2,512 genes, but computational limitations meant examining 288 ST131 genomes where only a single specimen per rST was examined across 1,230,995 SNPs from a 2.33 Mb core genome, with a larger set of 9,479 diverse *E. coli* [20]. Given that rST1503 alone may account for ~81% of ST131 and that outbreaks may comprise a single rST [21], our understanding of *E. coli* ST131 transmission dynamics and diversity within single STs may limit inferences of past, present and emerging MDR outbreaks. A deeper investigation of MDR ST131’s population structure, selective processes and ESBL gene evolution can illuminate its mechanisms of AMR, host colonisation and pathogenicity [10,14]. Exploring the evolutionary origins, transmission and spread of outbreaks requires extensive sampling to link variation at AMR genes with inferred adaptive and epidemiological patterns [22], and previous work suggests a high-resolution large-scale approach to bacterial epidemiological based on genomic data address these questions [23].

Deducing the evolutionary relationships based on the core genome permits the discovery of novel accessory genome events [24]. ST131 evolution has been punctuated by plasmid conjugation, plasmid recombination and mobile genetic element (MGE) rearrangements, particularly of the cefotaximase (CTX-M) class of ESBL genes,*bla*_CTX-M-14/15/27_ [25–27] that allow third-generation cephalosporin-resistance [28]. These *bla*_CTX-M_ gene changes correlate strongly with ST131 subclade differentiation, such that the most common one (C2) is typically *bla*_CTX-M-15_-positive [29]. ESBL and other virulence factor genes likely drive extraintestinal niche colonisation but vary across environments depending on MGE-driven mobility [10, 15, 29, 30]. When coupled with host immunity, this environmental niche effect results in negative frequency-dependent selection (NFDS) in the ST131 accessory genome, leading to a variable AMR gene repertoire [31] that has not yet to be explored within ST131’s subclades. In addition, applying an evolutionary pangenomic approach with core and accessory genome variation within subclades may inform on the origin of new genetic ST131 lineages.

Here, we aggregated all available ST131 Illumina HiSeq read libraries, and automated quality-control, genome *de novo* assembly, DNA read mapping and ESBL gene screening in the largest ST131 set examined thus far to reconstruct a core genome phylogeny and evaluated the epidemiology of clades and subclades. We established that the two most common C subclades (C1 and C2) co-circulated globally and that their ESBL gene composition was flexible. We hypothesise that the diversity of accessory genomes in isolates with near-identical core genomes due to ST131’s ability to retain newly acquired genes may be driven by environmental pressures.

## Results

### Collation, screening and generation of 4,071 high quality draft ST131 genome assemblies

We collated accession IDs and linked metadata for 4,071 high quality *de novo* genome assemblies whose DNA was isolated in 1967-2018 from diverse sources across 170 BioProjects (Supplementary Table S1) following thorough filtering steps (Fig. 1, see Methods). 4,070 genomes from Illumina HiSeq read libraries assembled using Unicycler had with N50s of 195,830 ± 57,037 bp (mean ± standard deviation), lengths of 5,136,890 ± 121,402 bp, 124.3 ± 74.8 contigs and 4,829 ± 142 genes (Supplementary Table S2). The sole PacBio assembly (AR_0058) had five contigs with a N50 of 4,923,470 bp and was 5,132,452 bp long.

**Figure 1.**
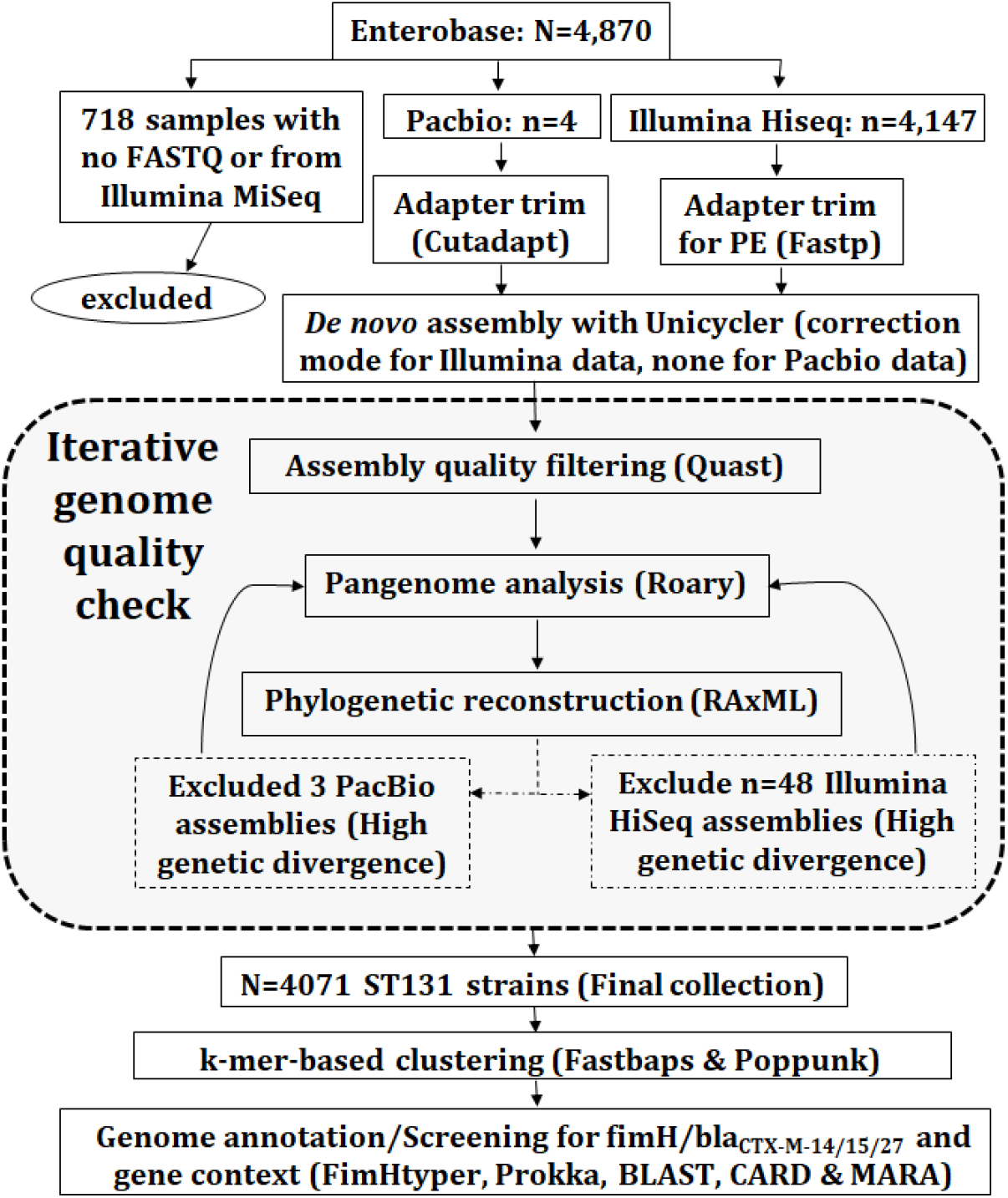
Methods summary: 4,870 read libraries were downloaded from Enterobase. 718 uninformative ones were excluded. Of those assessed, four were long read libraries (PacBio) and the rest were short paired-end reads (Illumina HiSeq). The adapters of the four PacBio and 4,147 Illumina reads were trimmed using Cutadapt and Fastp, respectively. The resulting adapter-free reads were assembled using Unicycler. An iterative genome quality check eliminated three PacBio and 77 Illumina libraries, yielding 4,071 as the final collection. Cleaned reads after Quast filtering were examined with Roary using Prokka annotation to evaluate the pangenomic diversity. Phylogenetic construction was performed by RAxML on the core genome. The assembled genomes were annotated and screened for AMR genes (including *bla*_CTX-M-14/15/27_) and their context. Genetically distinct clusters from the phylogeny were determined using Fastbaps. Distances between the core and accessory genomes of isolate pairs were estimated using Poppunk based on k-mer differences.

We assembled the pangenome of 4,071 ST131 isolates using NCTC13441 as a reference with Roary v3.11.2 [32] resulting in 26,479 genes, most of which were rare (Supplementary Fig. S1). *Bla*_CTX-M-15_-positive NCTC13441 was isolated in the UK in 2003, and was from subclade C2. 3,712 genes present in all isolates formed the core genome, and of 22,525 CDSs in the accessory one, 1,018 were shell genes in 15-95% of isolates and 21,507 (81% of the total) cloud genes in less than 15% of isolates (Supplementary Fig. S2). Cloud gene rates were a function of sample size, which explained most (r^2^=0.846, p=0.00012) of cloud gene number variation, but not that of core (r^2^=0.162), soft core (r^2^=0.258) or shell (r^2^=0.001) genes.

### Population structure classification shows three dominant ST131 C subclades

Clades A (n=414, 10.1%), Clade B (n=420, 10.3%) and Clade B0 (n=13, 0.3%) were relatively rare in comparison to the 3,224 in Clade C (79%) based on *fimH* typing. This showed 91% of Clade A had *fimH41*, 66% of Clade B had *fimH22*, 99% of Clade C had *fimH30*, and unexpectedly Clade B0 had *fimH30*, not *fimH27* (Table 1). Nine isolates were *fimH54*, of which eight were in Clade B [33].

**Table 1.** Number of ST131 in Clades A, B, B0, C0, C1 and C2. Isolates from Clade A mainly had *fimH*41 and were assigned to Fastbaps cluster 2. Clade B tended to have *fimH*22 as well as other *fimH* alleles, and were assigned to five Fastbaps groups (1/3/5/7/8). Clade C mainly had *fimH30* or *fimH*-like alleles, and were assigned to Fastbaps cluster 6 for C1 (aka C1_6), or clusters 4 (C2_4) or 9 (C2_9) for C2.

| Clade/<br>subclade | Fastbaps<br>Cluster IDs | *fimH* allele |  |  |  | Isolate<br>total |
| --- | --- | --- | --- | --- | --- | --- |
|  |  | 41 | 22 | 30 | Others |  |
| A | 2 | 376 |  |  | 38 | 414 |
| B | 1, 3, 5, 7, 8 | 8 | 277 |  | 135 | 420 |
| B0 | 8 |  |  | 13 |  | 13 |
| C0 | - |  |  | 51 | 1 | 52 |
| C1 | 6 |  |  | 1,111 | 10 | 1,121 |
| C2 | 4, 9 |  |  | 2,032 | 19 | 2,051 |
| Total |  | 384 | 277 | 3,207 | 203 | 4,071 |

Clustering of the 4,071 isolates based on 30,029 core genome SNPs with Fastbaps identified nine genetically distinct subclades (clusters 1-9) and two groups of heterogeneous or rare isolates (clusters 10 and 11) (Fig. 2). Clade A was mainly assigned to clusters 2 (n=407, 98.3%) and 11 (n=7, 2.7%) (Supplementary Fig. S3). Clade B isolates were in clusters 1 (n=90, 21.4%), 3 (n=96, 22.9%), 5 (n=64, 15.2%), 7 (n=115, 27.4%), 8 (n=4, 1.0%), 10 (n=34, 8.1%) and 11 (n=17, 4.0%). All isolates of Clade B0 were in cluster 8, suggesting that it was a subclade within Clade B. As a consequence of the heterogeneity of clusters 10 and 11, which members came from all three clades, these isolates were assumed to be unassigned to proper groups due to their rare number.

**Figure 2.**
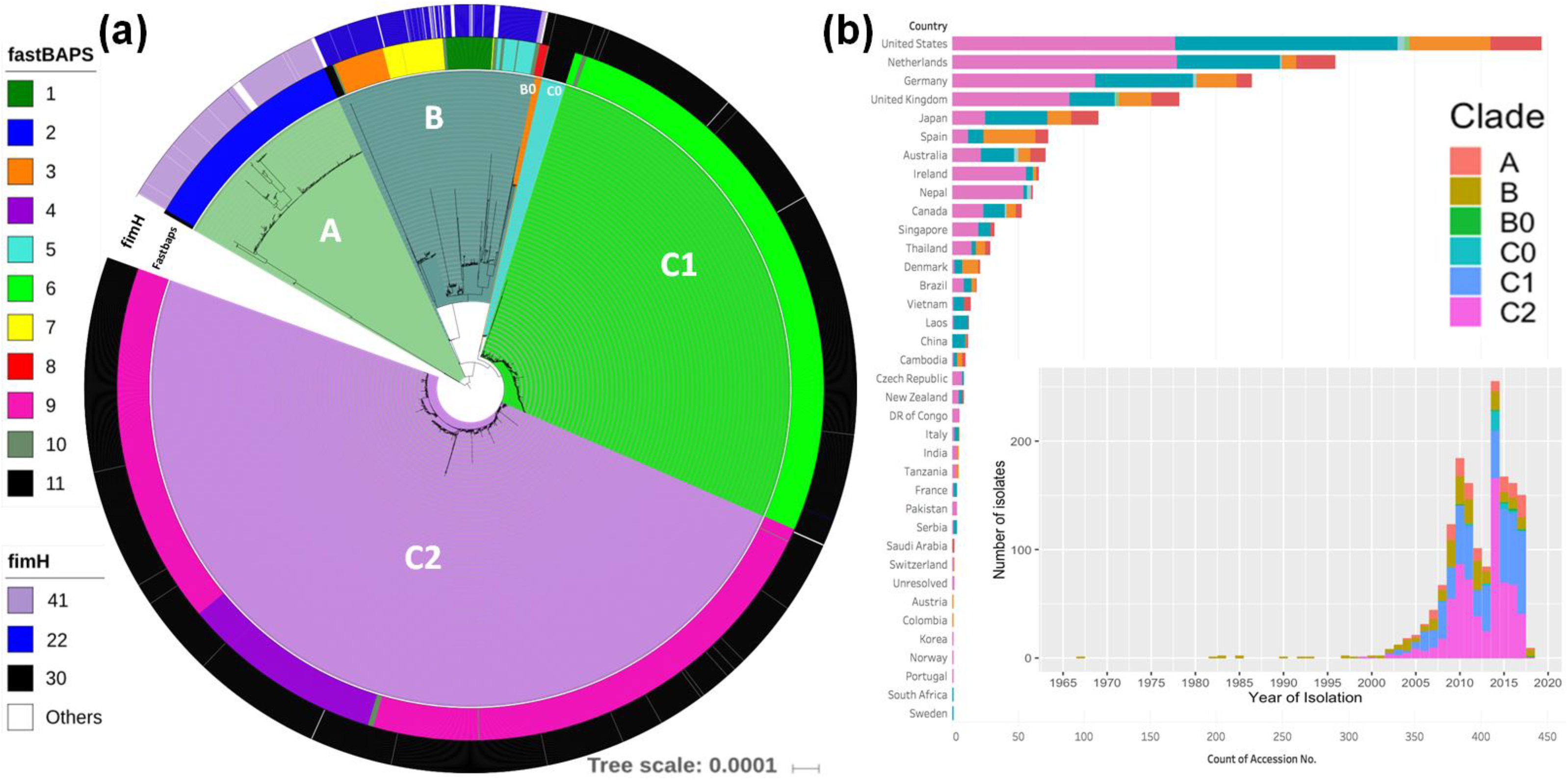
(a) A maximum likelihood phylogeny of 4,071 global ST131collection and (b) the distribution of these isolates across countries and over time. The phylogeny shows clades A (n=414 genomes, pale green), B (n=420 genomes, dark green), B0 (n=13 genomes, orange), C0 (n=52 genomes, blue), C1 (n=1,121 genomes, bright green) and C2 (n=2,051 genomes, purple). The phylogeny constructed with RAxML from the 30,029 chromosome-wide SNPs arising by mutation was visualized with iTol. The inner colored strip surrounding the tree represents the subgroups formed from Fastbaps clustering and the cluster (1-11) associated with each isolate. The outer colored strip surrounding the tree is the *fimH* allele (*H41* for Clade A (pink), *H22* for Clade B (blue), *H30* for Clade C (black) and other alleles (white). The histograms in (b) show the distribution of sampling across countries, and that out of the 4,071 ST131 genomes isolated from 1999 to 2018, 2,051 belong to C2 (pink), with most isolates coming from 2002-2017.

Clade C (n=3,224) had three main subclades determined by Fastbaps: C1_6, C2_4 and C2_9. C1 had 1,121 isolates: 1,113 isolates in Fastbaps cluster 6 (referred to as C1_6) with eight unassigned in cluster 10 (Fig. 3). C2 had 2,051 assemblies: 1,651 in cluster 9 (C2_9) and 386 in cluster 4 (C2_4). C0 (n=52 isolates) was mainly assigned to cluster 11, consistent with its heterogeneous nature [10]. 13 C2 genomes were assigned to cluster 10 and one to cluster 6.

**Figure 3.**
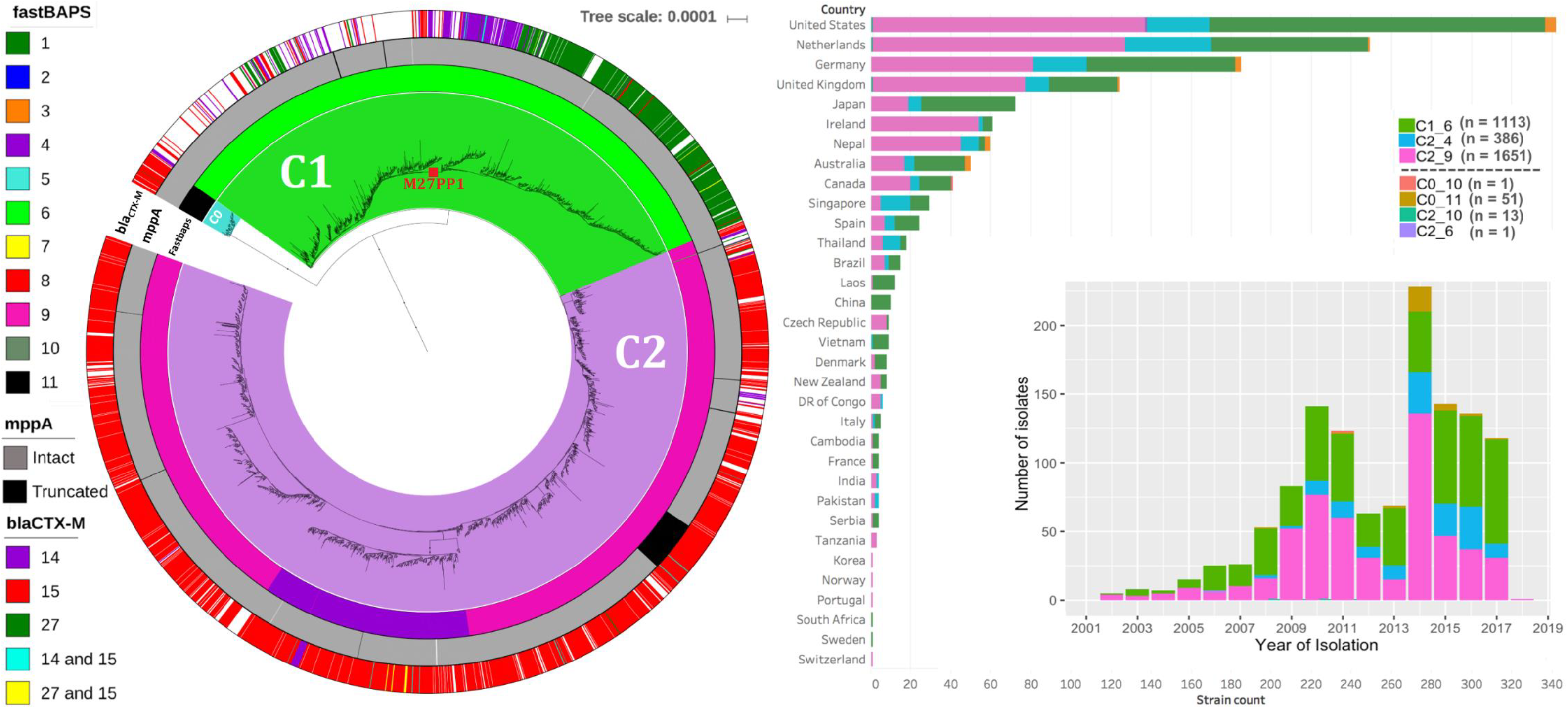
(a) A maximum likelihood phylogeny of 3,224 ST131 Clade C isolates and (b) the distribution of these isolates across countries and over time. The phylogeny shows C0 (n=52 genomes, blue), C1 (n=1,121 genomes, bright green) and C2 (n=2,051 genomes, purple). As per Fig. 2, the phylogeny constructed with RAxML from the 30,029 chromosome-wide SNPs arising by mutation was visualized with iTol. The inner colored strip surrounding the tree represents the subgroups formed from Fastbaps clustering and the cluster (1-11) associated with each isolate. This indicated most C0 were in Fastbaps cluster 11 (n=52 genomes) with a single isolate in cluster 10 (grey). Of the 1,121 C1 isolates, 1,113 formed Fastbaps cluster 6 (green) and eight were assigned cluster 10 (black). The C2 subclades corresponded to Fastbaps clusters 9 (C2_9, n=1,651 isolates, pink) and 4 (C2_4, n=386, dark purple). The middle colored strip indicates the isolates with an interrupted *mppA* gene (black) relative to the wild-type intact version (grey). The outer colored strip is the *bla*_CTX-M_ allele: 2,416 genomes had *bla*_CTX-M-14_, *bla*_CTX-M-15_, or *bla*_CTX-M-27_ genes, 1,790 genomes had *bla*_CTX-M-15_ (mainly C2), 177 genomes had *bla*_CTX-M-14_ (mainly C1) and 424 genomes had *bla*_CTX-M-27_ (mainly C1). The M27PP1 locus gain denoting the C1-M27 lineage (red box) was found in 468 C1_6 genomes (though independent events occurred too). The histograms in (b) show the distribution of sampling across countries, and that since 2002 both C1_6 (green) and C2_9 (pink) were common until the emergence of C2_4 (blue) and to a lesser extent C0 (brown).

### Epidemic subclades C1 and C2 co-circulate globally but with stable frequencies

NFDS in the accessory genome driven by AMR gene acquisition, ecological niche colonisation ability and host antigen recognition has stabilised the relative frequencies of ST131 and its clades over time relative to other STs [29,31]. Here, this pattern could be present for the clades A, B, C and three main C subclades, C1_6, C2_4 and C2_9 spanning 2002-2017 (1,596 out of 1,614 isolates that had year of isolation data) if their relative rates stabilised after emergence (Supplementary Fig. S4).

Of the 1,724 isolates with geographic information, 819 were from Europe, 499 North America, 294 Asia, 80 Oceania, 20 South America, and 12 Africa (Fig. 3) - the remaining 2,347 isolates (58%) had no geographic data (Supplementary Table S3). C1_6 was more common in North America (OR=1.57, 95% CI 1.25-1.96, p=0.0004) and rarer in Europe (OR=0.67, 95% CI 0.53-0.81, p=0.0004). C2_4 was more frequent in Asia (OR=1.75, 95% CI 1.18-2.56, p=0.019) and less so in North America (OR=0.61, 95% CI 0.40-0.91, p=0.042). Consequently, there was limited global population structure separating C1 and C2 isolates, suggesting that they have co-circulated globally for some time.

Based on common ancestry with Clade B, the origin of Clade C was in North America because Clade B isolates from 1967-1997 were solely isolated in the USA until one isolate in Spain in 1998. This fitted previous work timing the origin of C to 1985, *fimH*30 to 1986, and the FQ-R C1/C2 ancestor to 1991 [21] (or 1986 [29]), consistent with a North American source. However, the earliest C representative here from Norway in 1999 was a *bla*_CTX-M_-negative *bla*_TEM-1B_-positive FQ-R one from C2_9 (ERR1912633 [34]).

The origin of C2_4 was unclear: although the earliest isolate was in 2008 from the USA, the most basal branches within C2_4 had isolates spanning a range of continents, and C2_4’s long ancestral branch implied that it originated prior to 2008 (Supplementary Fig. S5). The isolates most closely related to C2_4 were a group of 11 assigned to C2_10 that had limited country and year of isolation data bar one from the UK in 2011, one from the USA in 2009, one from the Netherlands (Supplementary Fig. S3). The next most closely related group were nine C2_9 isolates that had no geographic data bar two from Australia in 2017.

### Variable *bla*_CTX-M-14/15/27_ gene prevalence across time, geography and ST131 subclades

Alignment of the 4,071 assemblies against *bla*_CTX-M-14/15/27_ genes and CARD with BLAST showed that these ESBL genes were more common in Clade C (75%) than Clade A (45%) than Clade B (4%) (Fig. 4). Few isolates were both *bla*_CTX-M-14/15_-positive (0.4%) or *bla*_CTX-M-15/27_-positive (0.3%) (Table 2). Of the 2,408 *bla*_CTX-M_-positive Clade C isolates, 1,782 isolates were *bla*_CTX-M-15_, 424 isolates were *bla*_CTX-M-27_, 177 isolates were *bla*_CTX-M-14_, 15 isolates were *bla*_CTX-M-14/15_, and 10 isolates were *bla*_CTX-M-15/27_ (Supplementary Fig. S6) such that the rates were highest in C2_4 (93.8%) followed by C0 (90%), C2_9 (82.6%) and C1_6 (57%) (Supplementary Fig. S7). The earliest *bla*_CTX-M_-positive Clade C genome was from Canada in 2000 (ERR161284, C2_9, [12]). 88% (339 of 386) of C2_4 isolates and 81% (1,338 of 1,651) of C2_9 isolates were *bla*_CTX-M-15_-positive with limited geographic or temporal structure (Fig. 4). This reiterated that the C2 ancestor was *bla*_CTX-M-15_-positive whose gains of other *bla*_CTX-M_ genes were likely rare local events.

**Table 2.** ST131 subclades’ *bla*_CTX-M-14/15/27_ genes. B0 (n=13) is not shown because it had no *bla*_CTX-M_ genes. The chromosomal gene *mppA* as intact or interrupted, where truncation of this gene was indicative of a chromosomal insertion of a *bla*_CTX-M-15_-positive TU.

| Subclade | Number | *bla*<sub>CTX-M</sub> allele numbers per isolate |  |  |  |  | *mppA* gene |  |
| --- | --- | --- | --- | --- | --- | --- | --- | --- |
|  |  | 14 | 14+15 | 15 | 15+27 | 27 | Interrupted | Intact |
| A | 414 | 65 | 1 | 66 | 2 | 51 |  | 414 |
| B | 420 | 7 | 1 | 9 |  | 1 | 11 | 409 |
| C0 | 52 |  |  | 46 |  | 1 |  | 52 |
| C1_6 | 1,113 | 149 | 6 | 59 | 3 | 418 | 3 | 1,108 |
| C2_4 | 386 | 12 | 3 | 339 | 7 | 1 | 1 | 382 |
| C2_9 | 1,651 | 16 | 6 | 1,338 |  | 4 | 90 | 1,561 |

**Figure 4.**
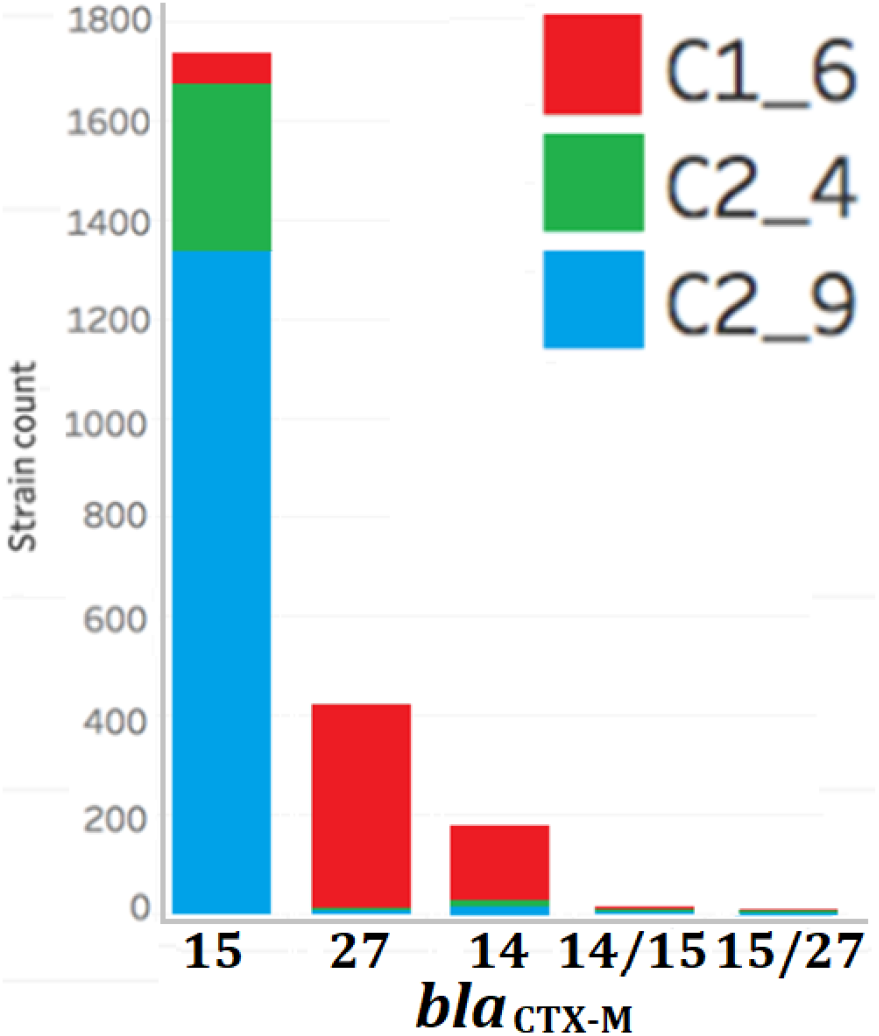
Frequencies of *bla*_CTX-M_ alleles in C subclades C1_6 (red), C2_4 (green) and C2_9 (blue).

C1_6 had a different *bla*_CTX-M_ gene rates to Clade C: *bla*_CTX-M-27_ (38%) was more common than *bla*_CTX-M-14_ (14%) or *bla*_CTX-M-15_ (6%) (Table 2). Of the *bla*_CTX-M_-positive isolates from subclade C1, the earliest was from 2002 (with a *bla*_CTX-M-14_-gene). The earliest *bla*_CTX-M-27_-positive and *bla*_CTX-M-15_-positive ones followed in 2004 and 2008, respectively. Subclade C1_6 was found in Japan only until detection in both China and Canada in 2005. *Bla*_CTX-M-14_ (but not *bla*_CTX-M-27_) was more common in Asia (OR=4.4, 95% CI 2.21-8.85, p=0.00007) as previously [35], and both *bla*_CTX-M-15_ and *bla*_CTX-M-27_ were global.

Previous work showed that C1 subclade C1-M27 with the prophage-like genomic island M27PP1 was a key driver of MDR ST131 [36]. Here, 468 (42%) of C1_6 isolates had M27PP1, and 97% of these were *bla*_CTX-M-27_-positive (397 of 410 with *bla*_CTX_ gene data) as expected (Supplementary Table S3) [36]. These formed a single clade (Fig. 3) that was globally disseminated since the earliest date for a C1-M27 isolate here was 2004 in Japan (DRR050997, [36]). 74 of these C1-M27 isolates subsequently acquired the related prophage-like genomic island M27PP2, most likely as a single ancestral event given their monophyly here. Five subclade C2 isolates and one from Clade A had M27PP1: all three of the C2_9 isolates with *bla*_CTX_ gene data were *bla*_CTX-M-27_-positive, and the earliest of these was from Japan in 2010 (DRR051016) [36].

### Diverse genomic locations of the *bla*_CTX-M-14/15/27_ genes’ contigs across ST131 subclades

Screening of the 505,761 contigs from the 4,071 assemblies for *bla*_CTX-M-14/15/27_-positive ones identified diverse structures and contexts based on annotation with MARA (Supplementary Table S4). 90 genomes from C2_9 had a *bla*_CTX-M-15_ gene in a transposition unit (TU) flanked by a 1,658 bp 5’ IS*Ecp1* and 3’ orf477 as a 2,971 bp IS*Ecp1*-*bla*_*CTX-M-15*_-orf477Δ TU followed by a 3’ 5.8 Kb Tn*2* (Supplementary Fig. S8, Supplementary Table S5), verified previously using long reads [26]. Normally this TU is on an IncF plasmid [35], but selected C2_9 assemblies had a chromosomal insertion of this TU at the *mppA* gene (Supplementary Fig. S9) as shown previously [21]. Here, 13 additional genomes with source information had this insertion, indicating that C2_9 had spread to Thailand (as in [14]), Singapore and the Democratic Republic of Congo by 2014 [37], consistent with a global spread. One C2_4 isolate from Pakistan in 2012 also had this TU inserted at *mppA* (SRR1610051 [38]), affirming that insertions at *mppA* recur due to local homology to IS*Ecp1*’s 3’ inverted repeat [39–40].

### Inter-clade but not intra-clade accessory genome divergence

Accessory genome composition varies across genetically distinct groups due to ecological niche specialisation [15]: this was supported here at the clade level across the 4,071 isolates by a positive correlation of pairwise core and accessory genome distances measured with Poppunk [23] (Fig. 5). This matched work on an *E. coli* dataset with n=218 ST131 [29], and was evident in a higher shell gene number in Clade B compared to Clades A and C (Table 3), suggesting higher diversity in Clade B (Supplementary Fig. S10).

**Table 3.** The pangenome composition of ST131 clades and subclades showed stable core, soft core and shell genomes with open pangenomes (*alpha*). The average *alpha* was determined for sample sizes from 30 to the maximum per group (see Fig. 6): for all ST131 this was 0.812±0.024 (mean ± standard deviation); for A 0.780±0.045; B 0.796±0.046; C 0.766±0.031; C1_6 0.749±0.017; C2_4 0.754±0.077; and C2_9 0.799±0.038. The groups’ cloud gene rates correlated with sample size following a power law model. B0 (n=13) was included with B. Eight C1_10 and 16 C2_6/C2_10 isolates not assigned to clear Fastbaps clusters were not examined.

| Clades | All | A | B | C | C1_6 | C2_4 | C2_9 |
| --- | --- | --- | --- | --- | --- | --- | --- |
| #isolates | 4,071 | 414 | 433 | 3,200 | 1,113 | 384 | 1,651 |
| *alpha* | 0.823 | 0.807 | 0.762 | 0.806 | 0.755 | 0.696 | 0.822 |
| Average *alpha* | 0.812 | 0.780 | 0.796 | 0.766 | 0.749 | 0.754 | 0.799 |
| Total genes | 26,479 | 12,163 | 16,323 | 21,304 | 16,490 | 10,322 | 15,485 |
| Core genes | 3,712 | 3,798 | 3,771 | 3,916 | 3,843 | 4,109 | 4,019 |
| Soft core genes | 242 | 292 | 281 | 334 | 424 | 380 | 354 |
| Shell genes | 1,018 | 764 | 1,437 | 731 | 571 | 642 | 566 |
| Cloud genes | 21,507 | 7,309 | 10,834 | 16,323 | 11,652 | 5,191 | 10,546 |

**Figure 5.**
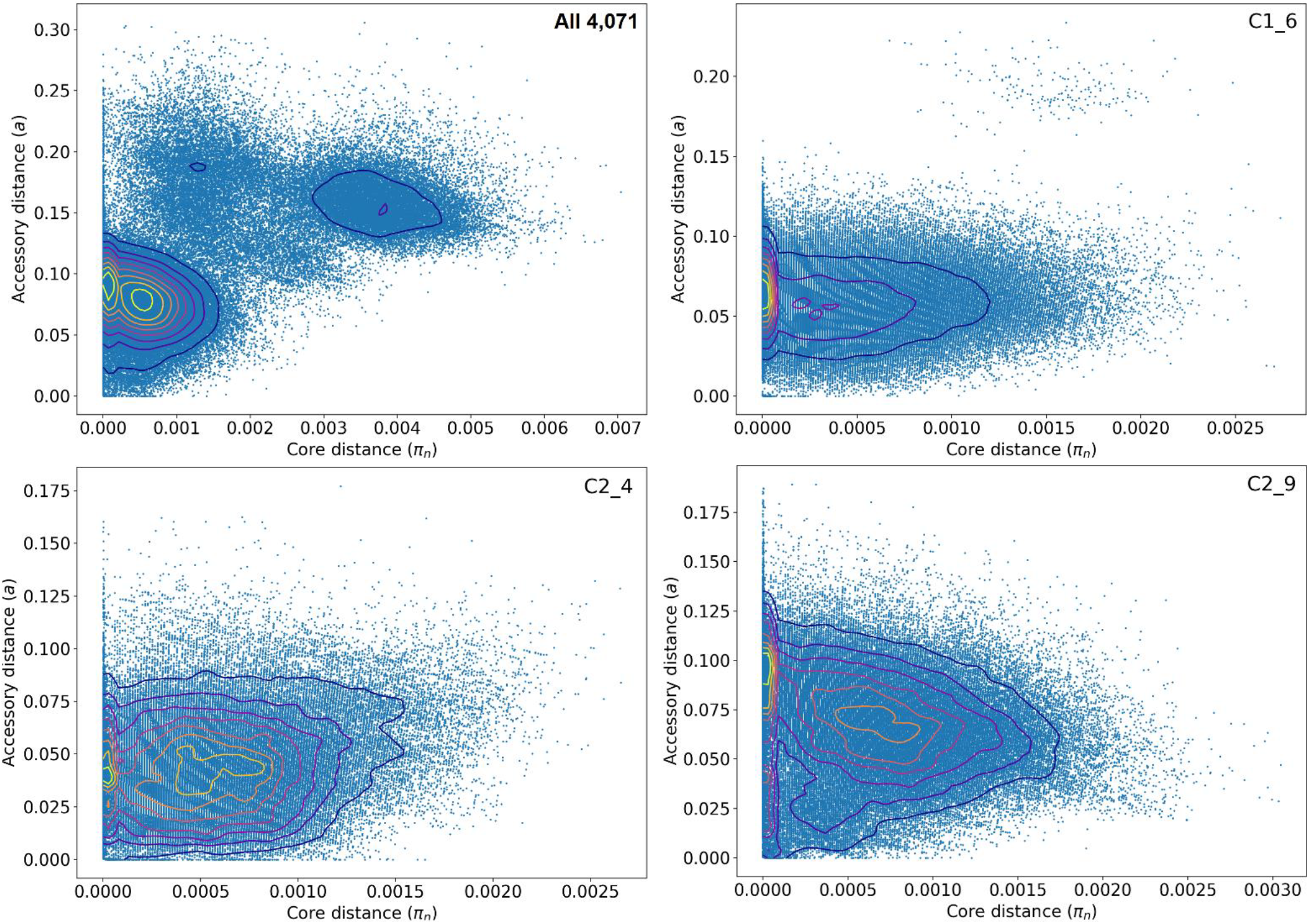
The distribution of core (*π*, x-axis) and accessory pairwise genome distances (*a*, y-axis) with blue dots indicating isolate pairs and the contours indicating dot density (higher in yellow). Top left: All 4,071 assemblies displayed pairwise differences such that the contours indicated the three main clades: Clade A at *π*=0.0038, *a*=0.15; Clade B at *π*=0.0014, *a*=0.18; Clade C at both *π*=0.0005, *a*=0.08 and *π*=0.0001, *a*=0.09. Top right: 1,113 subclade C1_6 assemblies had a peaks mainly at *π*≤0.001, *a*=0.06. Bottom left: 386 subclade C2_4 assemblies had peaks at *π*=0.0006, *a*=0.045 and *π*=0.0001, *a*=0.040. Bottom right: 1,651 subclade C2_9 assemblies had peaks at *π*=0.0007, *a*=0.065 and *π*=0.0, *a*=0.090. Results for 2,416 *bla*_CTX-M_-positive Clade C assemblies and 52 subclade C0 assemblies were similar. Within subclades C1_6, C2_4, C2_9, isolates had more diverse accessory genomes compared to their core ones.

Previous work on *E. coli* [41] employed a metric of pangenome openness (*alpha*) was similarly applied to our Roary pangenome results here to compare with previous findings that n=648 Clade C isolates had a marginally more open genome (smaller *alpha*) than n=140 from Clade B, and both were more open than n=70 Clade A isolates [31]. Although our initial results revealed more open pangenome (*alpha*) in Clade B (0.762) than Clade A (0.807) or Clade C (0.806) or the whole collection (0.823), *alpha* was higher for small (<250) sample sizes here (Fig. 6) as indicated before [41]. *Alpha* estimates averaged across the sample size placed Clade C as more open than Clade A or Clade B (Table 3), highlighting a partial dependence of *alpha* on sample size that was removed once the sample sizes >250 when the relative rate of new genes became constant (Supplementary Fig. S11). The average *alpha* for *250 ≤ n ≤ 386* showed Clade C (0.716) was more open than Clade B (0.753) than Clade A (0.795) or the whole collection (0.771) (Supplementary Fig. S12) The upper limit of n=386 was corresponded to the smallest group size, which was for C2_4.

**Figure 6.**
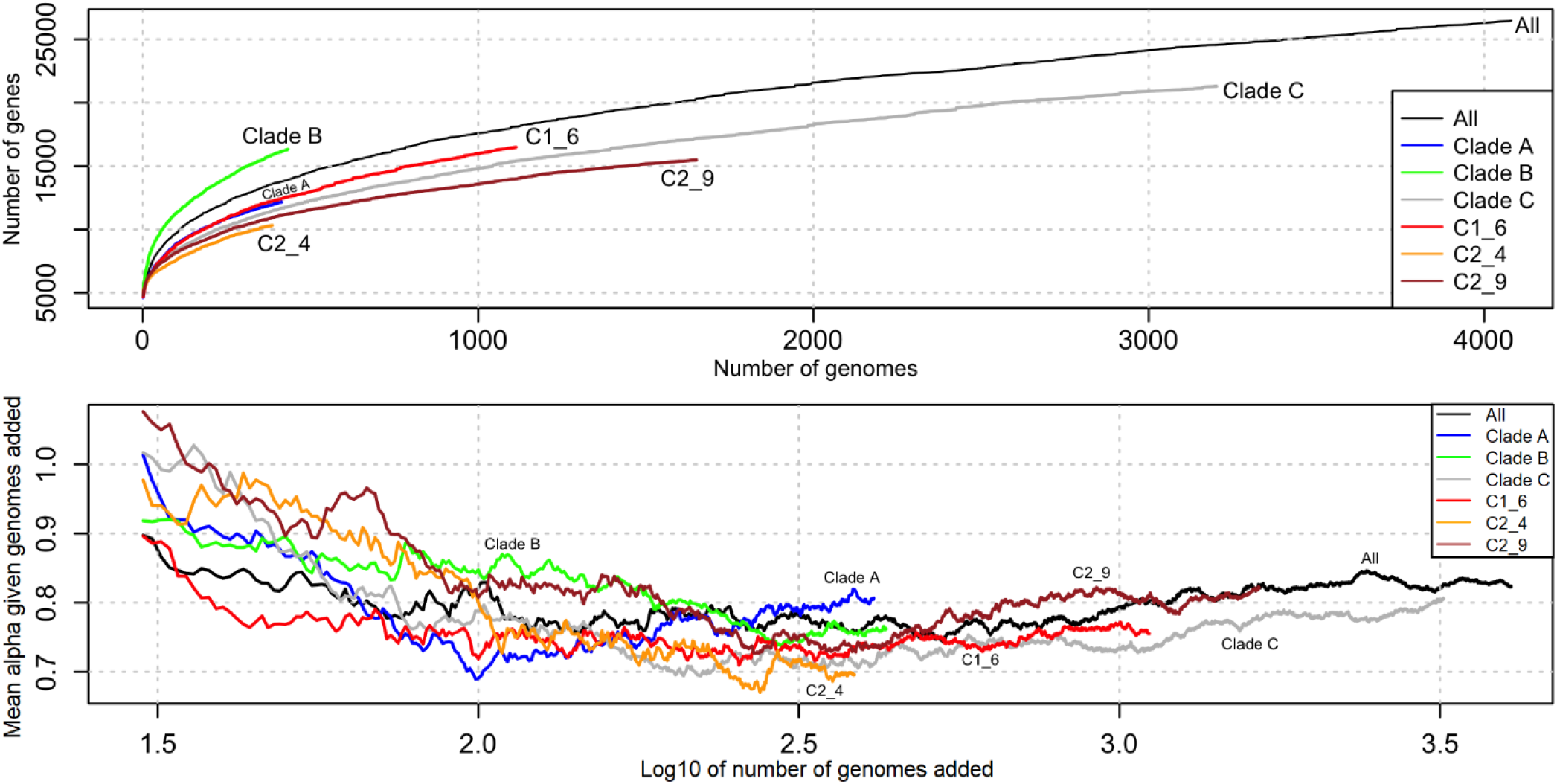
Top: The average number of genes in the ST131 pangenome (y-axis) increased as the 4,071 genomes were added (x-axis) indicating an open pangenome for the whole collection (black), as well as its clades and subclades: Clade A (blue), Clade B (green), Clade C (grey), subclade C1_6 (red), subclade C2_4 (orange) and subclade C2_9 (brown). Below: *Alpha* varied with numbers of genomes sampled (shown here for >30 genomes) and was more independent from sample number once the number of genomes examined about >250. Note that the x-axis’ log_10_ scale.

Within the C subclades, the pairwise core and accessory genome distances were not correlated: the accessory genomes varied extensively even with nearly identical core genomes (Fig. 5). Subclade C2_4 had a more open pangenome (0.696) than subclade C1_6 (0.755) or subclade C2_9 (0.822), which was evident when adjusting for the differing sample sizes (Fig. 6), though the average *alpha* placed C2_4 (0.754) and C1_6 (0.749) as about equally more open than C2_9 (0.799, Table 3). As above, the relative *alpha* levels were retained when *alpha* was averaged from >250 isolates up to the smallest (subclade C2_4) sample size with C2_4 (0.705) and C1_6 (0.731) as more open than C2_9 (0.742) (Supplementary Fig. S12).

Given that the high accessory genome diversity within subclades independent of core genome composition, the observed shell gene numbers were compared to expected values adjusted for sample number and gene frequency category change to investigate shell gene overlap across clades and subclades (see Methods). Pooled groups with divergent accessory genomes should have more net shell genes, whereas similar accessory genomes should have fewer shell genes. Clades B and C together had 6% less shell genes, whereas Clade A had an excess of 1% when combined with Clade C and 6% with Clade B (Supplementary Table S6). Within subclades, there was a small shell gene excess for C1_6 combined with C2_9 (3%), but C2_4’s shell gene composition differed from both C1_6 (22% excess) and C2_9 (23% excess, Supplementary Table S6). The same trend was observed for C2_4 combined with Clade A (41% excess) or Clade B (5%) in contrast to C1_6 (16% with Clade A, −8% with Clade B) and C2_9 (16% with Clade A, −11% with Clade B), indicative of more unique shell genes in subclade C2_4.

## Discussion

By collating all available ST131 genomes to produce 4,071 high-quality draft assemblies, we reconstructed their phylogenetic relationships to show that ST131 was dominated by subclades C1 and C2. For isolates with *bla*_CTX-M_ gene data, subclade C2 was 98% *bla*_CTX-M-15_-positive in contrast to C1 that had either *bla*_CTX-M-27_ (66%) or *bla*_CTX-M-14_ (24%) genes. Although the subclade C1 ancestor may have been *bla*_CTX-M-14_-positive, *bla*_CTX-M-27_’s increasing levels in C1 and its higher ceftazidime resistance due to a D240G substitution also in *bla*_CTX-M-15_ [42] indicated it will become more common. Although the subclades had different origins and ancestral ESBL gene compositions, both have become global with relatively consistent frequencies and minor differences in rates due to differing evolutionary patterns after emerging [14]. This worldwide co-circulation suggested newer lineages could become globally disseminated, with implications for infection control if they have altered host adhesion abilities (like *fimH30* [43]) or AMR variants (like FQ-R or *bla*_CTX-M-15_). This was highlighted by the emergence of C2_4, the C2_9 subgroup with an IS*Ecp1*-*bla*_CTX-M-15_-orf477-*Tn2* TU *mppA* chromosomal insertion, and many other contemporary examples such as *bla*_OXA-48_-producing ST131 [44]. Within C1, the *bla*_CTX-M-27_-positive C1-M27 lineage emerged in this study as an increasingly common cause of infection globally. Tracking plasmid, MGEs and ESBL genes must be a key component of disease monitoring to consider potential future bacterial outbreaks’ spectrum of AMR.

Horizontal DNA transfer allows *E. coli* adapt to new ecological niches and contributes to its dynamic accessory genome [45] where the cloud gene number increases with isolate number and diversity. Our analysis of this large collection’s core (3,712) and accessory (22,525) genes extended previous work showing that 283 predominantly ST131 isolates had 16,236 genes in an open pangenome with a core of 3,079 genes [46], 21% less than the core genome count here. Nonetheless, ST131’s accessory genome may be streamlined: a more genetically diverse set of 1,509 *E. coli* including 266 ST131 had a core genome of 1,744 genes and a 62,753 cloud genes [29], and an *E. coli-Shigella* core genome had 2,608 genes among a total of 128,193 genes [41].

NFDS posits that genes associated with adaptation to new hosts, antibiotics and competitors using the same resources remain at intermediate frequencies [31]. Pangenome openness and shell gene sharing across clades supported inter-clade structure resulting from ecological specialisation [47], with Clade A more different to Clades B and C. Within subclades, isolates with minimal core genome differences could have divergent accessory genomes, implying that plasmid, ESBL gene and MGE changes may be detected better using pangenomic approaches than assessing the core alone [48].

Global coordination of data processing and bioinformatic interpretation can help identify, trace and control disease outbreaks [49], for which resolving recent transmissions may be limited by sampling [50]. Expanding numbers of non-human isolates and the diversity of geographic regions sampled would help clarify potential sources of *E. coli*’s ESBL genes, for which there was no evidence of retail meat [51] or livestock [52] as reservoirs for blood stream infections thus far, though transfer of bacteria has occurred [53]. Better epidemiological information coupled with genome-sequencing [54] could allow inference of adaptations across lineages [55], such as *bla*_CTX-M-15_-positive ST8313, a putative descendant of ST131 subclade C2 [5].

## Methods

### Study selection and data extraction

Data on 4,870 *E. coli* ST131 genomes and linked metadata was collected using an automated text-mining algorithm using a Python implementation of Selenium (Selenium-python.readthedocs.io) from Enterobase (https://enterobase.warwick.ac.uk [56]) on the 10^th^ of September 2018 as previously described [57]. This was used to download read libraries the European Nucleotide Archive (ENA) (www.ebi.ac.uk/ena) [58] and NCBI Short Read Archive (SRA) databases as FASTQ files, restricted to complete libraries not labelled as “traces” (Fig. 1). Of the initial 4,870 read libraries, 4,264 were paired-end (PE) Illumina HiSeq ones and four were PacBio, in addition to the PacBio-sequenced NCTC13441 genome used as a reference in this study. 495 libraries predominantly from Illumina MiSeq platforms were not examined to avoid platform-specific artefacts.

### Illumina HiSeq read data quality control, trimming and correction

Of the above 4,264 PE Illumina HiSeq read libraries, 4,147 passed stringent quality control. This was implemented using Fastp v0.12.3 [59] to trim sequencing adapters, remove reads with low base quality scores (phred score <30) or ambiguous (N) bases, correct mismatched base pairs in overlapped regions and cut poly-G tracts at 3’ ends (Supplementary Table S2). Individual bases in reads were corrected by BayesHammer in SPAdes v3.11 [60]. Quality control metrics were examined at each step: across the whole collection as a batch report using MultiQC v1.4 [61] and on individual FASTQ files using FastQC v0.11.8 (www.bioinformatics.babraham.ac.uk/projects/fastqc/). 117 (2.7%) Illumina HiSeq libraries did not pass quality control.

### Illumina HiSeq read library genome assembly

The 4,147 Illumina HiSeq libraries passing quality control were *de novo* assembled using Unicycler v4.6 in bold mode to merge contigs where possible [62]. This used SPAdes v3.12 [63] to generate an initial assembly polished by Pilon v1.22 [64], which ran iteratively until no further corrections were required by the optimal assembly. This approach was similar to Enterobase’s [56], though Enterobase used BBMap in BBTools [65], SPAdes v3.10 and BWA [66] during assembly [20].

### Reference PacBio genome quality control and assembly

The ST131 reference genome NCTC13441 was isolated in the UK in 2003 and was in subclade C2 [67]. It had a 5,174,631 bp chromosome with 4,983 protein-coding genes and one pEK499-like type IncFIA/FIIA plasmid with two *bla*_CTX-M-15_ gene copies (accession ERS530440). Although four further PacBio read libraries were initially included to test genome assembly contiguity and ESBL gene context using longer read libraries, only one passed assembly annotation screening (AR_0058, accession SRR5749732 [38]). Its adapters were removed using Cutadapt v1.18 [68] followed by excluding duplicate reads with Unicycler v0.4.6. Base correction was implemented during genome assembly with Unicycler via SPAdes v3.12, and the genome assembly was iteratively polished by Racon v1.3.1 until no further corrections were required [69]. This 5,132,452 bp assembly had five contigs and 5,506 genes was assigned to C1 and had no IS*Ecp1*.

### Quality checking and annotation identifies 4,071 genome assemblies for investigation

For the 4,147 Illumina HiSeq assemblies and single PacBio assembly, quality was verified with Quast v5.0 [70] based on the N50, numbers of predicted genes and open-reading frames, and numbers of contigs with mis-assemblies. The quality of the short read *de novo* assemblies was comparable to previous work whose requirements required assembly length in the range 3.7-6.4 Mb with <800 contigs and <5% low-quality sites [20]. Initial annotation of 4,147 Illumina HiSeq assemblies using Prokka v1.10 [71] suggested 77 assemblies had a distinct gene composition and should be excluded because they were either genetically divergent, did not assemble adequately, or had sub-standard read libraries. As a result, 4,070 Illumina HiSeq genome assemblies were selected (Supplementary Table S2) and aligned against the reference genome NCTC13441 and PacBio assembly AR_0058 (Supplementary Table S3). These assemblies had lengths in the range 4.3-6.1 Mb with a mean and standard deviation of 5,137 ± 121 Kb. This identified 4,829 genes on average per assembly (range 3,942 to 5,749, Supplementary Fig. S1). The variation in numbers of genes per assembly was largely explained by the total assembly length (r^2^=0.959). 53% of the 4,071 had no source data and 12% of the remainder had a non-human source.

### Pangenome analysis to identify the core and accessory genomes

We created a pangenome based on the 4,072 annotation files using Roary v3.11.2 [32] with a 100% BLAST v2.6.0 identity threshold using the MAFFT v7.310 setting [72]. The resulting concatenated core CDS alignment spanning 1,244,619 bases and 3,712 genes scaffolded using NCTC13441 was used for core genome analyses. 242 soft core genes were found that may have been due to assembly errors or other artefacts. Pangenomes for each clade, C subclade and various combinations were also created for accessory genome evaluation.

### Phylogenetic reconstruction, population structure and subclade assignment

A maximum likelihood phylogeny was generated based on the core genome alignment of 4,071 genome assemblies with NCTC13441 as a reference across 30,029 SNPs (with 26,946 alignment patterns) for 50 iterations of RAxML v8.2.11 with a GTR model and gamma substitution rate heterogeneity [73]. 88% (3,585) of the assemblies were genetically unique. The total execution time on an Ubuntu v16.04 computer server with 256 Gb RAM using 52 threads was 24.4 days. Phylogenies were drawn and annotated using iTol v4.3.2 [74].

Clade classifications were initially based on published ST131 *fimH* phylogenetic analyses associating clade A with *fimH41*, B with *fimH22*, B0 with *fimH27*, and C with *fimH30* [21]. To classify the large number of isolates in the C subclades, we clustered the 30,029 core genome SNPs as a sparse matrix using a hierarchical Bayesian clustering algorithm implemented in Fastbaps v1.0 (Fast Hierarchical Bayesian Analysis of Population Structure [75]) in R v3.5.3 with packages ape v5.3, ggplot2 v3.1.1, ggtree v1.14.6 [76], maps v3.3.0 and phytools v6.60. This used default parameters except for a Dirichlet prior variance of 0.006.

The C1-M27 lineage was identified based on the prophage-like 11,894 bp M27PP1 region using the 3’ end of accession LC209430 [36], which was largely intact in the isolates with this element. The 19,352 bp M27PP2 prophage-like region was also examined using the 5’ end of the same accession. The monophyletic nature of the C1-M27 group was verified by examining their phylogenetic proximity, which contrasted with the paraphyletic M27PP2-positive C1 isolates, as well as other isolates from Clade A and C2 that were diverse.

### ESBL gene screening and contig visualisation

We screened for contigs with *bla*_CTX-M-14/15/27_ genes across the 4,071 assemblies’ 505,761 contigs using BLASTn [77] alignment of these three genes individually, and the Comprehensive Antibiotic Resistance Database (CARD) v3.0 requiring 100% identity for any match with a contig. Selected *bla*_CTX-M-14/15/27_-positive contigs were visualised using the Multiple Antibiotic Resistance Annotator (MARA) [78], R v3.5.2 and EasyFig v2.2.2 [79] to examine the local contig, MGE and gene annotation. A minority of isolates had incomplete contigs due to the small contig lengths. Frequencies of ST131 clades, subclades and their *bla*_CTX-M-14/15/27_ genes across geographic regions and time were examined with R packages dplyr v8.0.1, forcats v0.4.0, ggplot2 v3.1.1, ggridges v5.1, grid v3.5.2, plotly v4.9.0, plyr v1.8.4, purr v0.3.2, questionr v0.7.0, readr v1.3.1, rentrez v1.2.1, stringr v1.4.0, tibble v2.1.1, tidyr v0.8.3, tidyverse v1.2.1 and XML v3.98-1.19.

### Pangenome analysis to find shared and unique accessory genomes

Roary assigned genes to the core (*c*) and the accessory genomes, including the soft core (*s*), shell (*a*) and cloud (*d*). The expected shell gene number (*E*[*a*_*p*_]) from the Roary output for a given pooled set of isolates (*p*) taken from groups *i*=1..*k* was determined based on the shell gene number of group *i* (*a*_*i*_) weighted by the sample size (*n*_*i*_) corrected for the deficit in core (*c*_*i*_) and soft core (*s*_*i*_) gene numbers: 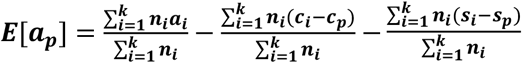. The excess fraction of shell genes observed was (*a*_*p*_ − *E*[*a*_*p*_])/*E*[*a*_*p*_]. Similarly, the expected cloud gene number *E*[*d*_*p*_] was computed from the cloud gene number of group *i* (*d*_*i*_) weighted by the sample size (*n*_*i*_) adjusted for the difference in core (*c*_*i*_), soft core (*s*_*i*_) and shell (*a*_*i*_) gene numbers: 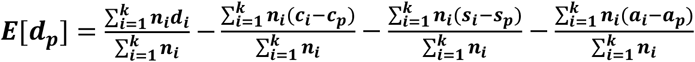. The excess fraction of cloud genes observed was (*d*_*p*_ − *E*[*d*_*p*_])/*E*[*d*_*p*_].

Pangenome openness (*alpha*) was quantified from Roary results as *Δn* = *kN*^*-alpha*^ where Δ*n* was the number of new genes across *N* genome assemblies with *n* genes in total [41] with R packages poweRlaw v0.70.2, igraph v1.2.4.1 and VGAM v1.1.1 (Supplementary Fig. S11). This power-law regression approximated Heaps’ law (from *n* = *ĸN*^*gamma*^ for *alpha=1-gamma*) such that an open pangenome has *alpha* < 1 and a closed one *alpha* > 1 [80]. Previously, diverse *E. coli* had *alpha = 0.625* where *alpha* had a partial negative correlation with *N* [41]. Similarly, *alpha* was ~0.877 for ST131 clade C, ~0.898 for B, ~0.958 for A, and ~0.951 for all ST131, suggesting *alpha* was higher when genetically distinct clades were combined [31].

### Accessory genome composition across clades and subclades

The relative pairwise genetic distances of the core (*π*) and accessory (*a*) genomes were compared for each clade, each C subclade and all *bla*_CTX-M_-positive clade C isolates using Poppunk (Population Partitioning Using Nucleotide Kmers), which can distinguish closely related genomes [23]. Poppunk used variable length pangenome k-mer comparisons with Mash v2.1 [81] and a Gaussian mixture model to examine the correlation of π and *a* per sample pair. This annotation- and alignment-free strategy complemented the approaches of Fastbaps, RAxML and Roary.

### Availability of materials and data

All raw sequence data (reads and/or assembled genomes) for the *E. coli* genomes analysed in this publication are in Supplementary Table S3, with their Bioproject IDs and associated study DOIs in Supplementary Table S1. The genome assemblies of the 4,071 *E. coli* ST131 are on Zenodo at https://zenodo.org/record/3341533 for 2,948 and at https://zenodo.org/record/3357944 for the remaining 1,123 files. The 4,071 *E. coli* ST131 genome annotation files are on Zenodo at https://zenodo.org/record/3341535 for 4,069 and at https://zenodo.org/record/3357914 for the final two files. An interactive version of the phylogeny generated by Poppunk for the 4,071 ST131 assemblies is on MicroReact at https://microreact.org/project/oD6K_fL2d - this includes a Newick tree file for download.

## Acknowledgements

We thank Marius Kinderis (Dublin City University, Ireland) for assistance in implementing the text mining algorithm. This work was funded by a DCU O’Hare Ph.D. fellowship and a DCU Enhancing Performance grant.

## Authors’ contributions

AD and TD organised funding, designed the study, performed genomic analyses and wrote the paper. AD led text mining, bioinformatic processing, data visualisation and phylogenetic analyses. All authors read and approved the final manuscript.

## Supplementary Figures

**Supplementary Figure S1.** Annotation of the 4,071 ST131 genomes (along with NCTC13441) using Prokka (a) identified 4,829 genes on average per assembly with a minimum of 3,942 and maximum of 5,749. (b) Of 26,479 gene clusters detected using Roary, 3,712 comprised the core genome (blue) spanning 1,244,619 bases based on pangenome analysis with 242 soft core genes (yellow), 1,018 shell genes (navy) and 21,507 cloud genes (light blue).

**Supplementary Figure S2.** Pangenome analysis of the effect of increasing the number of genomes (x-axis) showed (left) that few new genes were discovered (black line), but that the number of unique genes associated with the cloud gene set increased consistently (dashed line). (Right) The core genome composition across all 4,071 assemblies was stable once >200 genomes were included (“Conserved genes”, solid line), whereas the total number of genes increased without plateauing (dashed line). There was a median of 2.1 additional genes per additional isolate in this collection.

**Supplementary Figure S3.** Hierarchical sub-clustering of 4,071 isolates using Fastbaps based on 30,029 SNPs. Clusters are indicated by numerical numbers in bold red on above the blue bars. There were nine major clusters found and two (clusters 10 and 11) were dispersed among the collection.

**Supplementary Figure S4.** The annual fraction of isolates from (top) C1_6 (red), C2_4 (green) and C2_9 (blue) and (bottom) clades A (red), B (beige), B0 (green), C0 (light blue), C1 (dark blue) and C2 (mauve) showed consistent levels with no clear evidence of periodic radiation and fixation of new lineages.

**Supplementary Figure S5.** Phylogeny of 382 C2_4 isolates rooted using C2_9 isolates (not shown). The first C2_4 isolate found was in 2008 in the USA, but C2_4’s long ancestral branch imply that it arose several years prior to this. 90% (349) of C2_4 isolates had a *bla*_*CTX-M-15*_ gene. The outer ring shows isolates’ continents of origin, with Africa in orange (from the Democratic Republic of Congo); Asia in dark green (India n=1, Japan n=6, Nepal n=9, Pakistan n=2, Singapore n=15, Thailand n=9); Europe in bright green (Germany n=27, Ireland n=2, Italy n=1, the Netherlands n=43, Spain n=5, UK n=12, Vietnam n=1); Oceania in yellow (Australia n=5); and South America in light blue (both from Brazil). Four additional C2_4 isolates (n=386 in total) with long branches are not shown for clarity.

**Supplementary Figure S6.** Detection of at least one C1_6 (red), C2_4 (green) or C2_9 (blue) isolate per country (x-axis) per year from 2000-2019 (y-axis).

**Supplementary Figure S7.** Bubble chart of C1_6, C2_4 and C2_9 colored by their *bla*_*CTX-M*_ alleles (*bla*_*CTX-M-14*_ in orange, *bla*_*CTX-M-15*_ in turquoise, *bla*_*CTX-M-27*_ in green, and *bla*_*CTX-M-14/15*_ together in red) with country of origin shown within each bubble (where known). The area of each bubble corresponds with the relative frequency of that particular combination of subclade, country and *bla*_*CTX-M*_ allele.

**Supplementary Figure S8.** Representative examples of the *bla*_CTX-M-14/15/27_-positive contigs’ ESBL genes and MGEs annotated using Prokka and MARA. Some contigs were too short to show additional annotations, which can be estimated based on the longer contigs. (a) C1_6 had the most *bla*_CTX-M-14_-positive contigs that were typically in IS*Ecp1*-*bla*_*CTX-M-14*_-IS*903B* TUs, though with variations such as a 3’ IS*903C* element instead. (b) C2 tended to have *bla*_CTX-M-15_ flanked by a 5’ IS*Ecp1* and a Tn2 or orf-477-Tn2 at the 3’ as a 2,971 bp IS*Ecp1*-*bla*_*CTX-M-15*_-orf477Δ-*Tn2* TU. These were most likely on an IncF plasmid for the plasmid-encoded variants, but many within C2_9 had this TU inserted at the chromosomal *mppA* gene due to local sequence homology with IS*Ecp1*’s 14 bp 3’ inverted repeat (IRR). (c) C1_6 had the highest incidence of *bla*_CTX-M-27_-positive contigs that usually had a similar IS*Ecp1*-*bla*_*CTX-M-14*_-IS*903B* structure as shown in (a).

**Supplementary Figure S9.** Chromosomal insertion of *bla*_*CTX-M-15*_ indicated by an interrupted *mppA* gene was observed most commonly in C2. Intact *mppA* genes are shown in blue bars while the truncated ones were in red and were mainly observed in recent (2003-2017) C2_9 isolates.

**Supplementary Figure S10.** A phylogeny of all ST131 isolates (left) with the corresponding pangenome-wide gene presence (blue) and absence (white) frequencies per isolate represented for each of the 26,479 genes discovered. In the latter matrix, the 3,712 core genes are shown first on the left side, followed by the 242 soft core genes, 1,018 shell genes in 15-95% of isolates, and 21,507 cloud genes in <15% of isolates. Clades B (top and bottom clusters separated by green lines) and A (second from bottom separated from C by a red line) had core genome differences compared to C (middle bounded by green and red lines).

**Supplementary Figure S11.** Across the 4,071 genome assemblies (x-axis on a log10 scale), (top plot) the regression slope alpha (blue line) was estimated as 0.8231 such that the median number of new genes (y-axis) added per isolate (middle plot) was 2.1 (green points) and (bottom plot) the relative rate of new genes (red points) became constant once the number of isolates was >250 (or 10^2.4^). The rate of new genes (green points) and rate of change of new genes (red points) were fitted by loess curves with a span of 0.1 and a degree of two on the average number of new genes per isolate generated by Roary results.

**Supplementary Figure S12.** The variation in the number of genes and average alpha (y-axes) versus the numbers of genomes sampled (shown here for 251 to 357 genomes sampled, excluding missing data) (x-axis). Top: The number of genes added increased gradually across groups with the numbers of genomes, though with higher net diversity of genes in Clade B (green), next C1_6 (red), then Clade A (blue), followed by Clade C (grey), then by C2_9 (brown), and lastly C2_4 (orange). Bottom: The average alpha estimated varied with numbers of genomes showed a consistently more open genome for C2_4 (orange, average alpha = 0.705), followed by C (grey, average alpha = 0.716), then C1_6 (red, average alpha = 0.731), before C2_9 (brown, average alpha = 0.742), next B (green, average alpha = 0.753), the whole collection (black, average alpha = 0.771) and A (blue, average alpha = 0.795).

## Supplementary Tables

**Supplementary Table S1.** The 170 BioProject accession numbers with study and source information for the 4,071 ST131 genomes examined.

**Supplementary Table S2.** Quality statistics of the 8,140 Illumina read libraries and single PacBio read library associated with the 4,071 assemblies, including the proportion of duplicate reads, average GC content, mean sequence length (bp) and total number of sequences (millions) per library.

**Supplementary Table S3.** Metadata of 4,071 high quality ST131 genomes assessed. This includes the accession numbers, strain name, Bioproject accession numbers, *fimH* allele, clade assignment, Fastbaps cluster, subclade, *bla*_CTX-M_ allele, ISEcp1 count, year of isolation, city of isolation, country of isolation, continent of isolation, source niche type, M27PP1 presence, M27PP2 presence, total assembly length (bp), assembly N50, number of contigs in assembly, largest contig length (bp), GC content (%) and the number of genes in the assembly.

**Supplementary Table S4.** Selected (n=28) isolates whose *bla*_CTX-M_-positive contigs were visualised with MARA, including three from C0, nine from C1, two from C2_10, six from C2_4, and eight from C2_9.

**Supplementary Table S5.** The contig identifiers of the 2,623 ST131 assemblies with IS*Ecp1* elements along with the IS*Ecp1* lengths (maximum 1,655 bp).

**Supplementary Table S6.** Pangenome analysis for the collection and associated subsets, including the numbers of core (*c*), soft core (*s*), shell (*a*) and cloud (*d*) genes; the deficit in core, soft core and shell genes; the expected number of shell genes (*E[a]*) and percentage excess shell genes (*(a-E[a])/E[a]*); the expected number of cloud genes (*E[d]*) and percentage excess cloud genes (*(d-E[d])/E[d]*); and pangenome openness (*alpha*).

## List of abbreviations

AMR: Antimicrobial resistance
CARD: Comprehensive Antibiotic Resistance Database
cgMLST: Core genome MLST
CTX-M: Cefotaximase
*E. coli*: Escherichia coli
ENA: European Nucleotide Archive
ESBL: Extended-spectrum beta-lactamase
Fastbaps: Fast Hierarchical Bayesian Analysis of Population Structure
FQ-R: Fluoroquinolone-resistant
Inc: Incompatibility group
MARA: Multiple Antibiotic Resistance Annotator
MDR: Multidrug-resistant
MGE: Mobile genetic element
MLST: Multi-locus sequence typing
NCBI: National Centre for Biotechnology Information
NFDS: Negative frequency-dependent selection
PE: Paired-end
Poppunk: Population Partitioning Using Nucleotide Kmers
rST: Ribosomal MLST
SRA: Short Read Archive
ST: Sequence type
TU: Transposition unit
SNP: Single Nucleotide Polymorphism

