## Supplementary material for "An *Escherichia coli* ST131 pangenome atlas reveals population structure and evolution across 4,071 isolates": Suppl_Figures

### Supplementary Data for: An *Escherichia coli* ST131 pangenome atlas shows population structure and evolution across 4,071 isolates

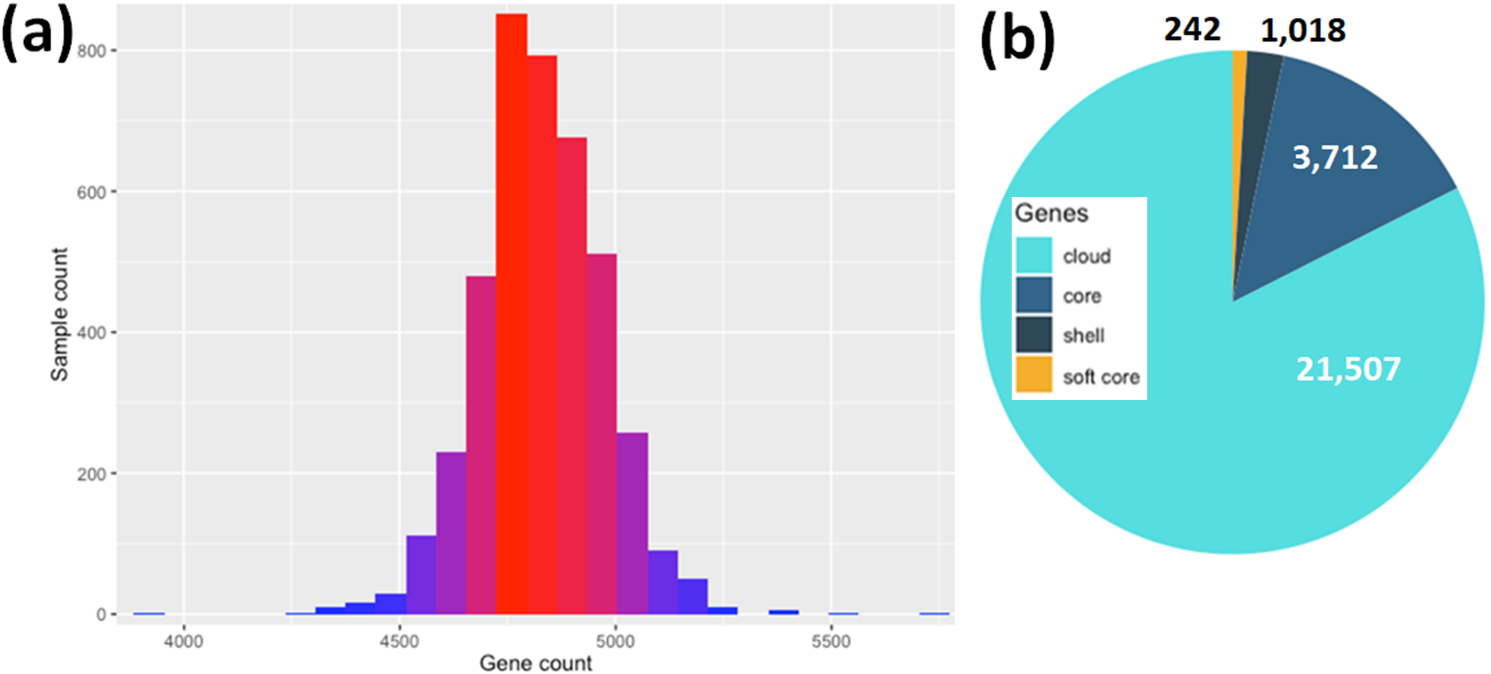

**Supplementary Fig. S1.** Annotation of the 4,071 ST131 genomes (along with NCTC13441) using Prokka (a) identified 4,829 genes on average per assembly with a minimum of 3,942 and maximum of 5,749. (b) Of 26,479 gene clusters detected using Roary, 3,712 comprised the core genome (blue) spanning 1,244,619 bases based on pangenome analysis with 242 soft core genes (yellow), 1,018 shell genes (navy) and 21,507 cloud genes (light blue).

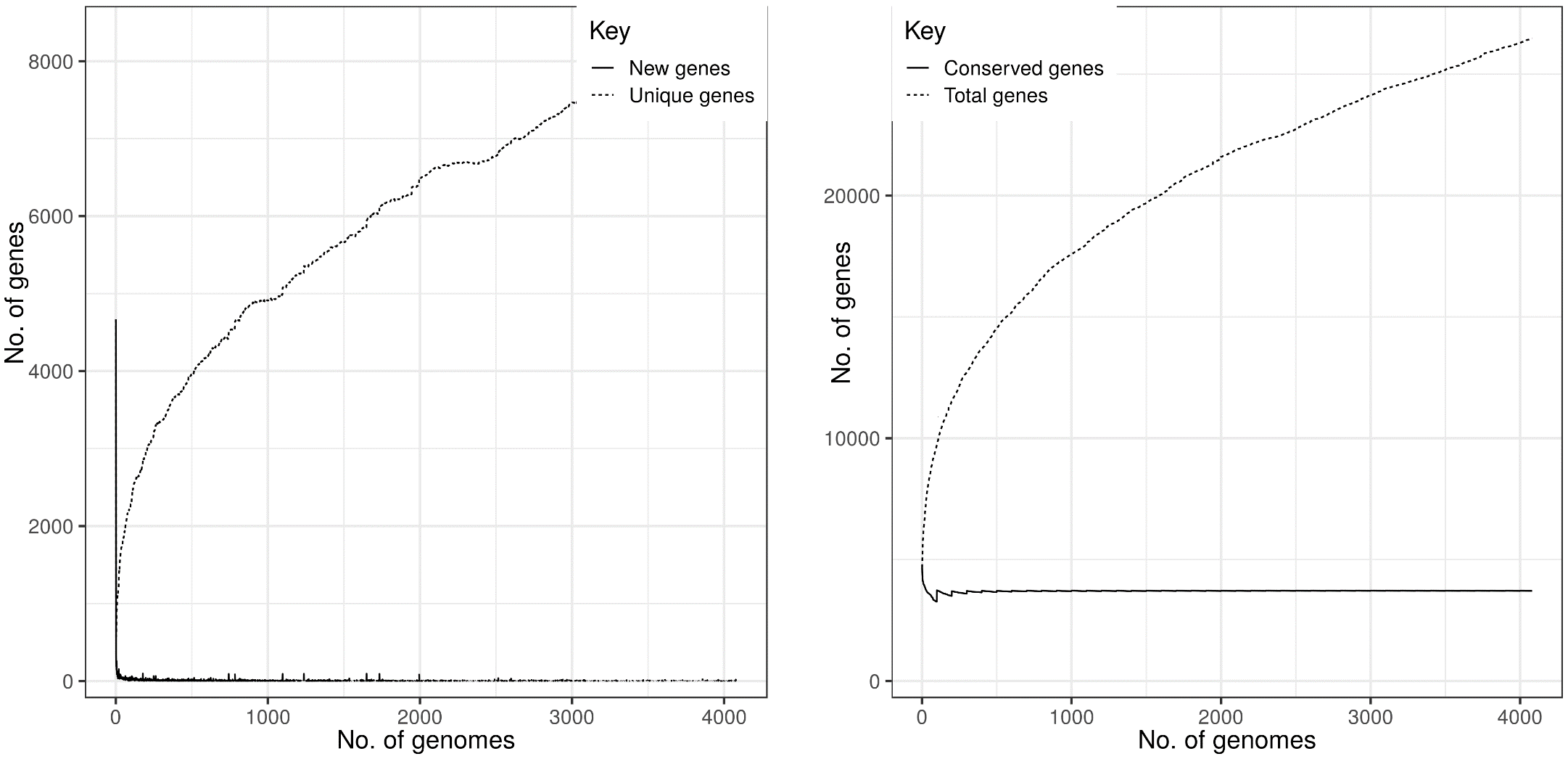

**Supplementary Fig. S2.** Pangenome analysis of the effect of increasing the number of genomes (x-axis) showed (left) that few new genes were discovered (black line), but that the number of unique genes associated with the cloud gene set increased consistently (dashed line). (Right) The core genome composition across all 4,071 assemblies was stable once >200 genomes were included (“Conserved genes”, solid line), whereas the total number of genes increased without plateauing (dashed line). There was a median of 2.1 additional genes per additional isolate in this collection.

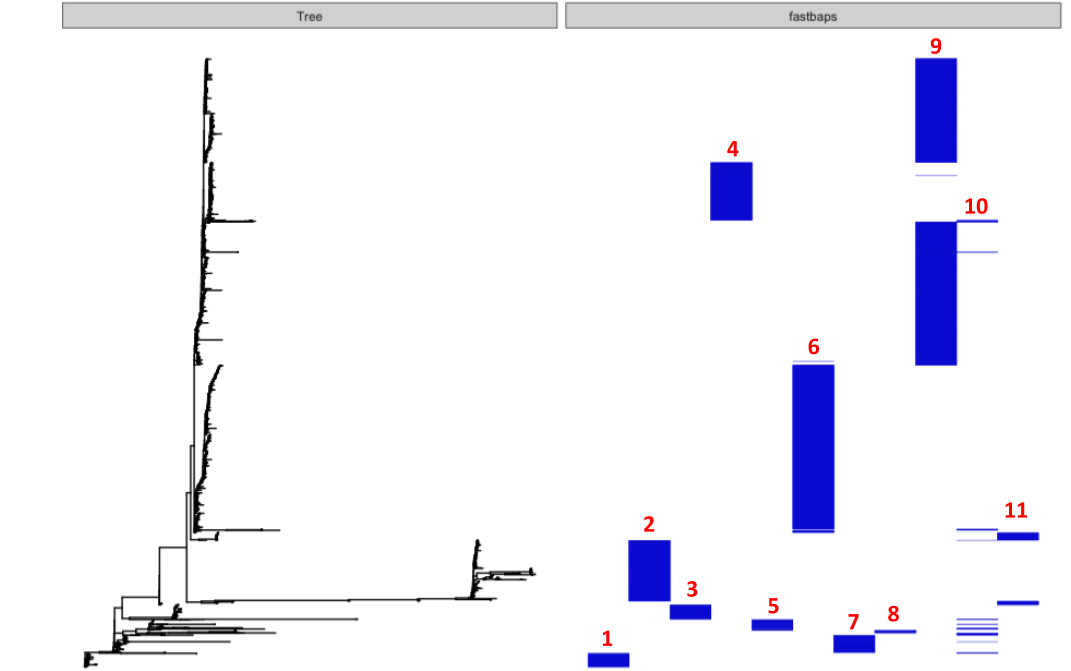

**Supplementary Fig. S3.** Hierarchical sub-clustering of 4,071 strains using Fastbaps based on 30,029 SNPs. Clusters are indicated by numerical numbers in bold red on above the blue bars. There were nine major clusters found and two (clusters 10 and 11) were dispersed among the collection.

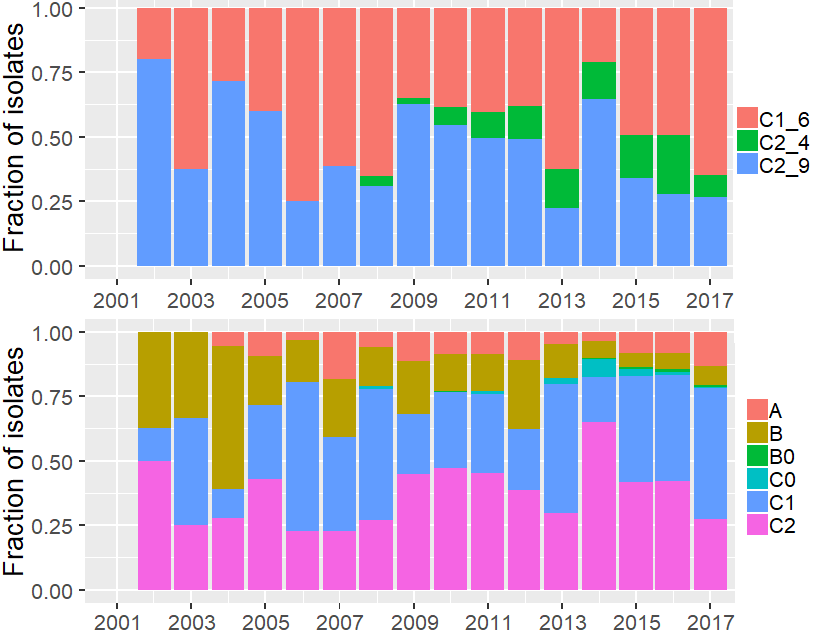

**Supplementary Fig. S4.** The annual fraction of samples from (top) C1_6 (red), C2_4 (green) and C2_9 (blue) and (bottom) clades A (red), B (beige), B0 (green), C0 (light blue), C1 (dark blue) and C2 (mauve) showed consistent levels with no clear evidence of periodic radiation and fixation of new lineages.

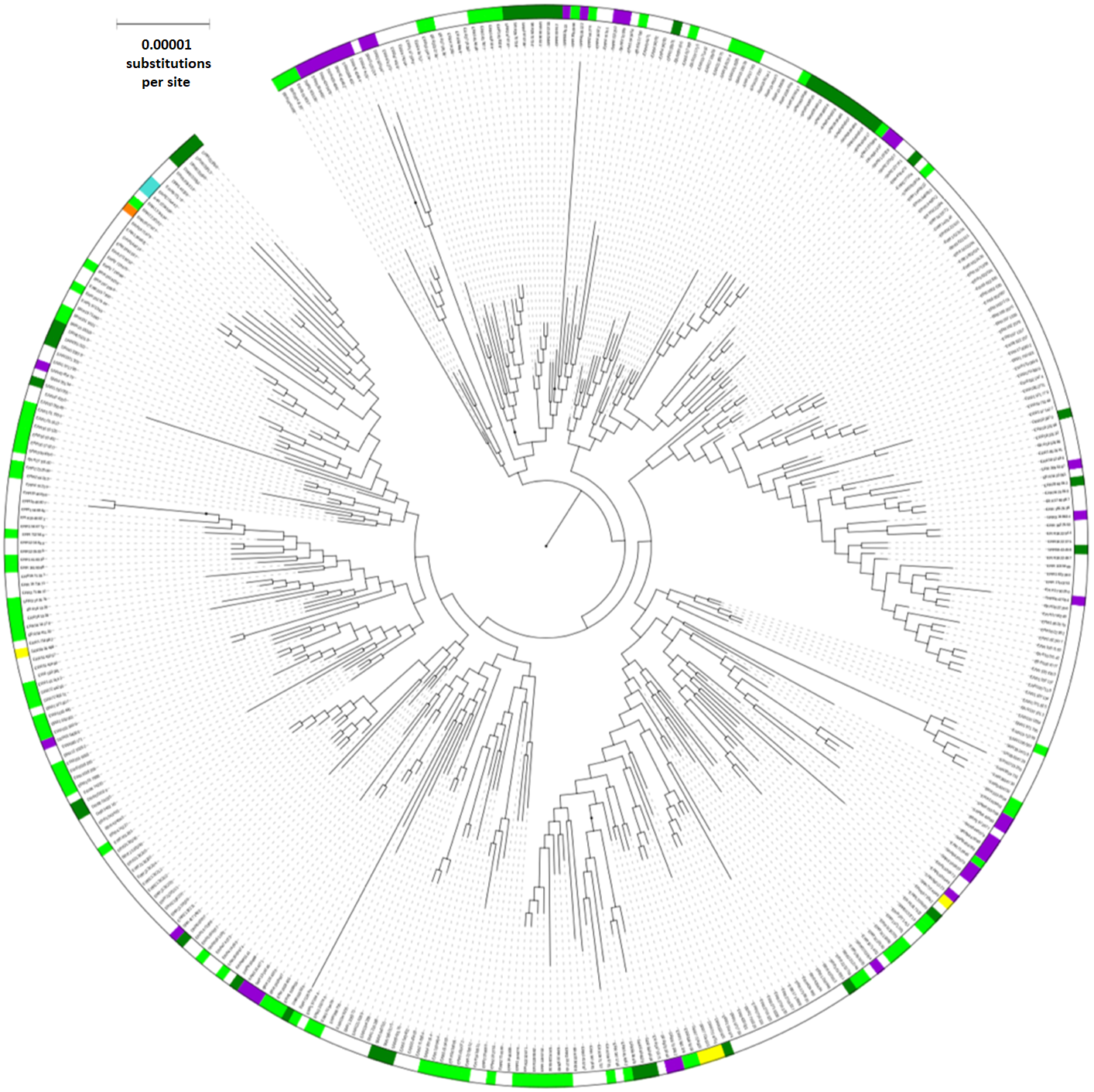
**Supplementary Fig. S5.** Phylogeny of 382 C2_4 strains rooted using C2_9 isolates (not shown). The first C2_4 isolate found was in 2008 in the USA, but C2_4’s long ancestral branch imply that it arose several years prior to this. 90% (349) of C2_4 isolates had a *bla_CTX-M_*_-15_ gene. The outer ring shows isolates’ continents of origin, with North America in purple (USA n=31, Canada n=4); Africa in orange (the Democratic Republic of Congo n=1); Asia in dark green (India n=1, Japan n=6, Nepal n=9, Pakistan n=2, Singapore n=15, Thailand n=9); Europe in bright green (Germany n=27, Ireland n=2, Italy n=1, the Netherlands n=43, Spain n=5, UK n=12, Vietnam n=1); Oceania in yellow (Australia n=5); and South America in light blue (both from Brazil). Four additional C2_4 isolates (n=386 in total) with long branches are not shown for clarity.

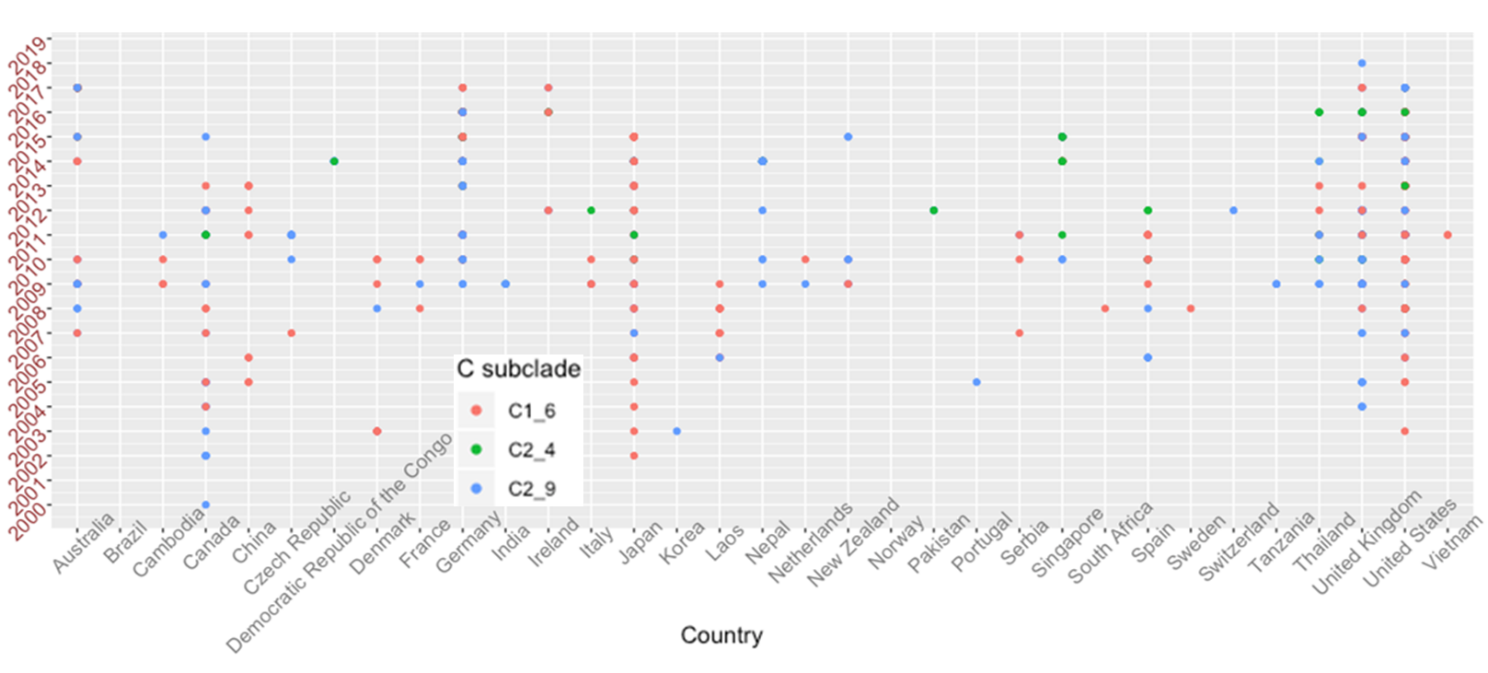

**Supplementary Fig. S6**. Discovery of at least one C1_6 (red), C2_4 (green) or C2_9 (blue) isolate per country (x-axis) per year from 2000-2019 (y-axis).

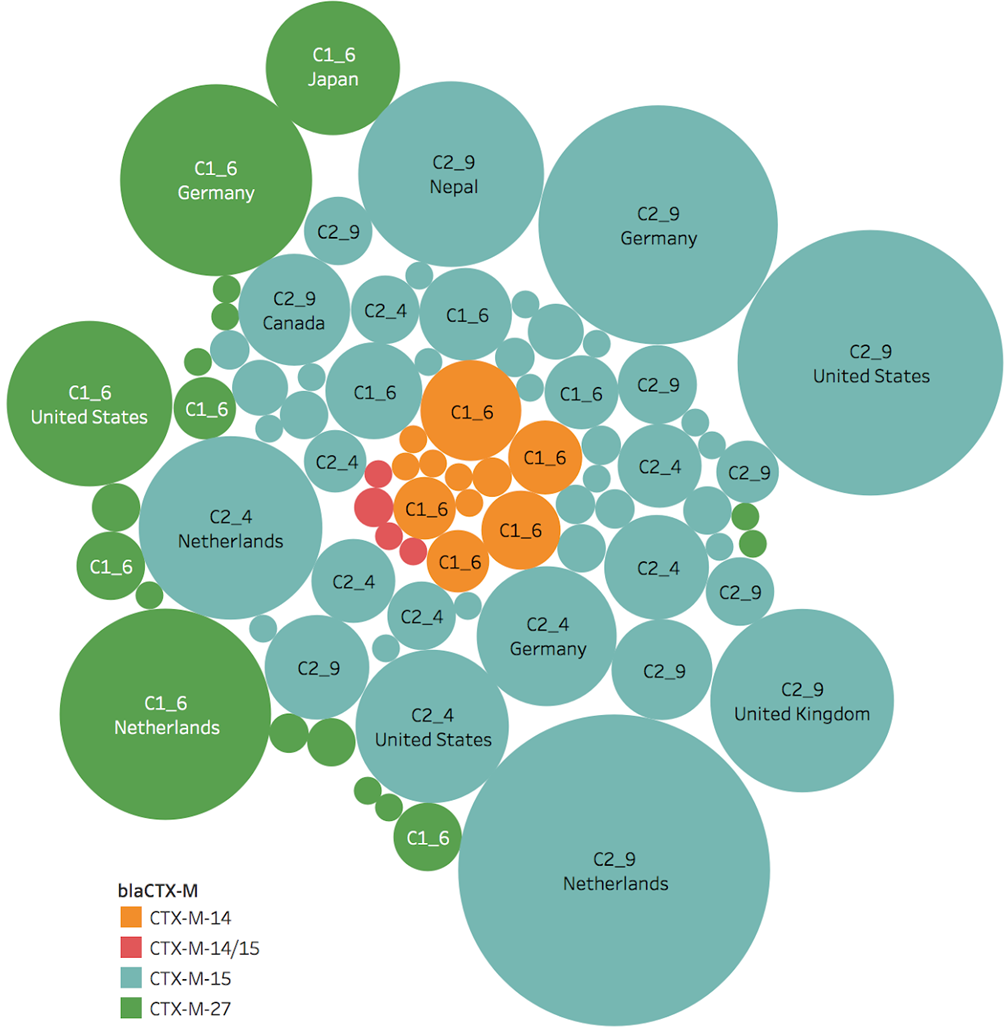
**Supplementary Fig. S7.** Bubble graph of C1_6, C2_4 and C2_9 colored by their *bla_CTX-M_* alleles (*bla_CTX-M-14_* in orange, *bla_CTX-M_*_-15_ in turquoise, *bla_CTX-M_*_-27_ in green, and *bla_CTX-M-14/15_* together in red) with the country of origin shown within each bubble (where known). The area of each bubble corresponds with the relative frequency of that particular combination of subclade, country and *bla_CTX-M_* allele. Only the countries with the higher levels of detected and published *bla_CTX-M_*-positive ST131 genomes are shown for visual clarity.

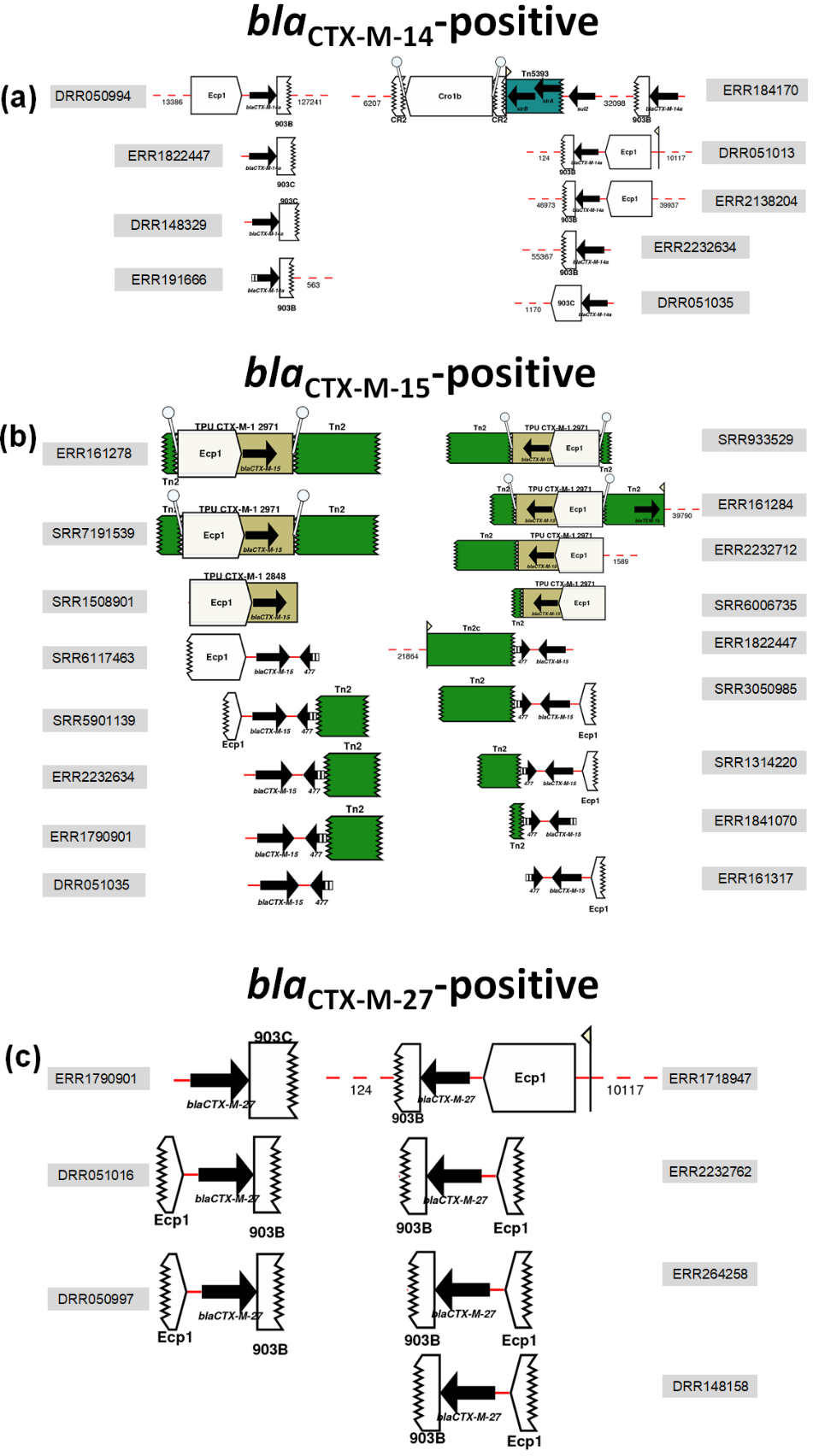
**Supplementary Fig. S8.** Representative examples of the *bla*_CTX-M-14/15/27_-positive contigs’ ESBL genes and MGEs annotated using Prokka and MARA. Some contigs were too short to show additional annotations, which can be estimated based on the longer contigs. (a) C1_6 had the most frequent incidence of *bla*_CTX-M-14_-positive contigs that were typically in IS*Ecp1*-*bla_CTX-M-14_*-IS*903B* TUs, though with variations such as a 3’ IS*903C* element instead. (b) C2 tended to have a *bla*_CTX-M-15_ gene flanked by a 5’ IS*Ecp1* and a Tn2 or the orf-477-Tn2 in tandem at the 3’ as a 2,971 bp IS*Ecp1*-*bla_CTX-M-15_*-orf477Δ-*Tn2* TU. These were most likely on an IncF plasmid for the plasmid-encoded variants, but many within C2_9 had this TU chromosomally inserted at the *mppA* gene due to local sequence homology with IS*Ecp1*’s 14-bp 3’ inverted repeat (IRR). (c) C1_6 had the highest incidence of *bla*_CTX-M-27_-positive contigs that typically had a similar IS*Ecp1*-*bla_CTX-M-14_*-IS*903B* structure as shown in (a).

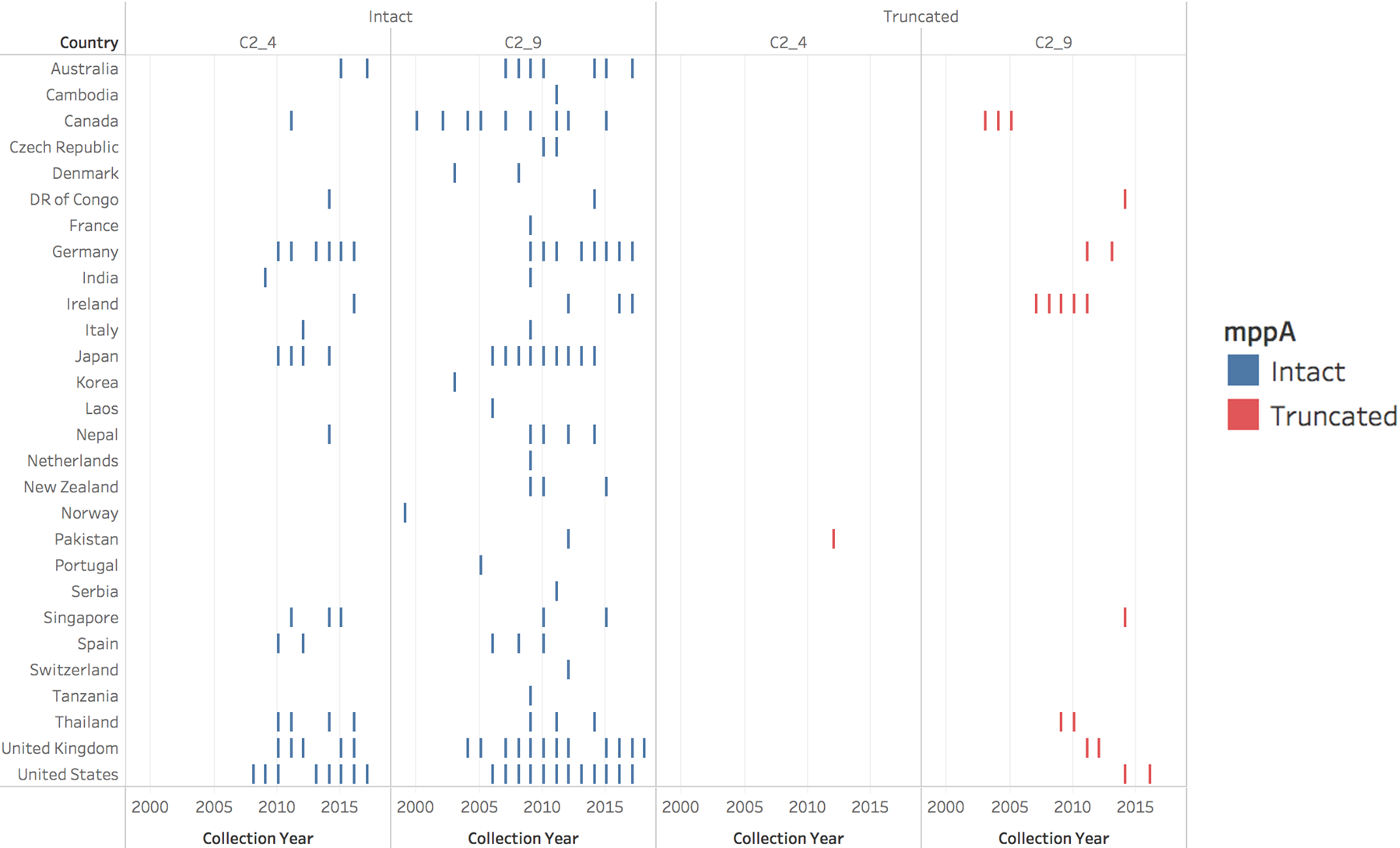

**Supplementary Fig. S9.** Chromosomal insertion of *bla_CTX-M-15_* indicated by a split *mppA* gene was observed most commonly in C2. Intact *mppA* genes are shown in blue bars while the truncated ones were in red and were mainly observed in recent (2003-2017) C2_9 samples.

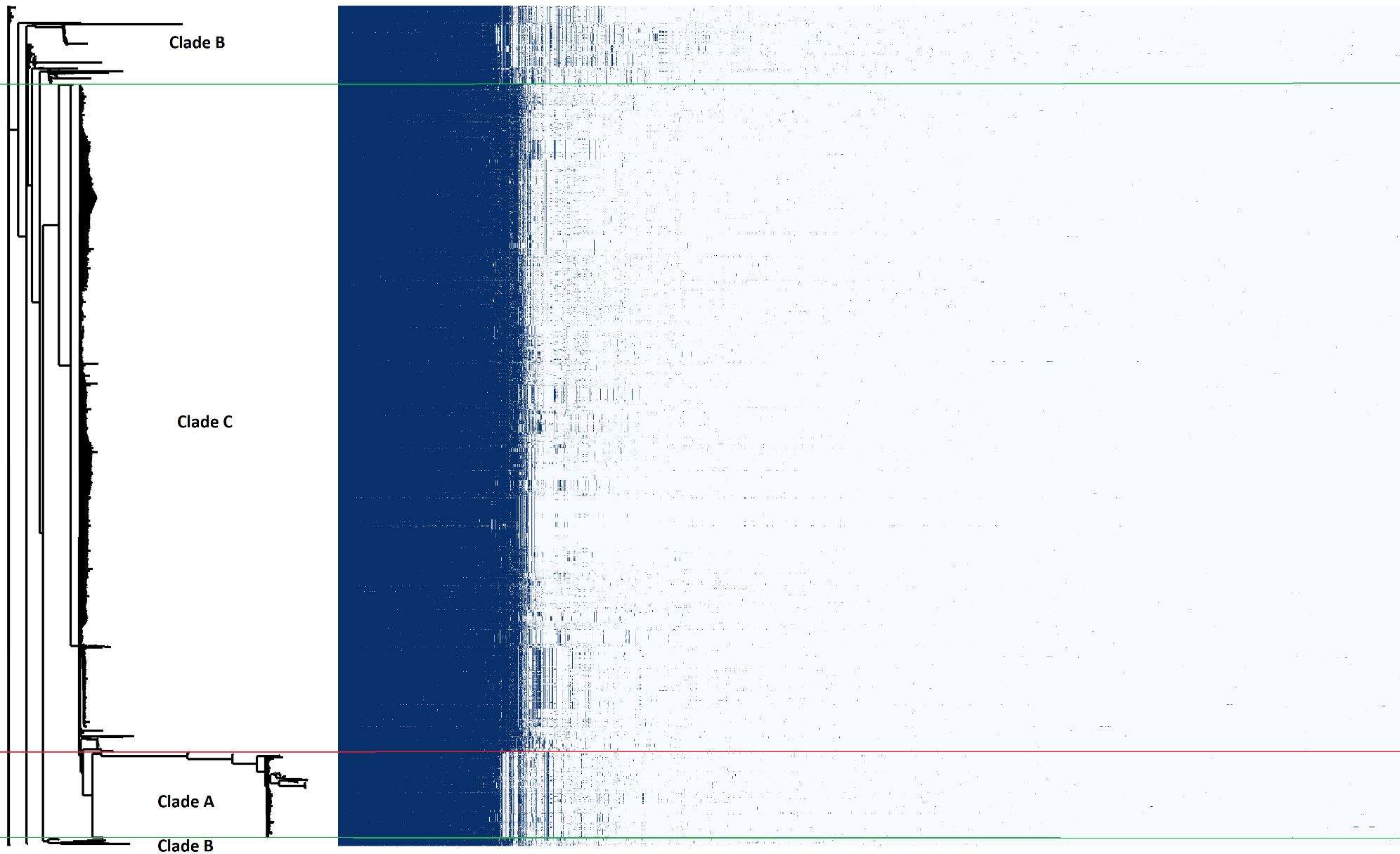

**Supplementary Fig. S10.** A phylogeny of all ST131 (left) with the corresponding pangenome-wide gene presence (blue) and absence (white) frequencies per isolate represented for each of the 26,479 genes discovered. In the latter matrix, the 3,712 core genes are shown first on the left side, followed by the 242 soft core genes, 1,018 shell genes in 15-95% of samples, and 21,507 cloud genes in <15% of samples. Clades B (top and bottom clusters separated by green lines) and A (2^nd^ from bottom separated from C by a red line) had core genome differences compared to C (middle bounded by green and red lines).

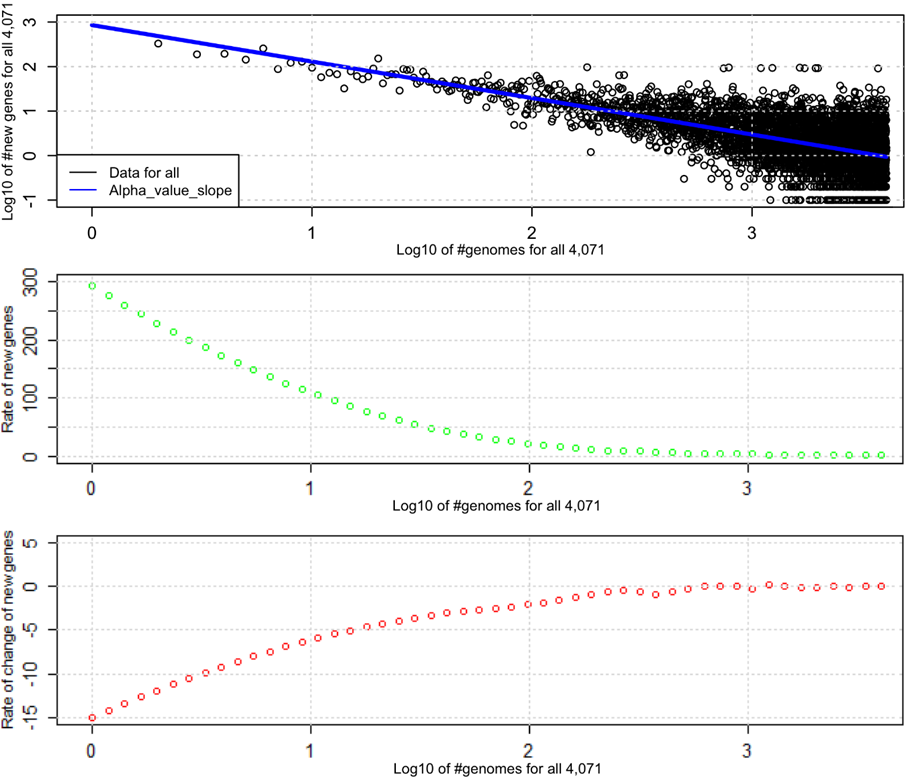

**Supplementary Fig. S11.** Across the 4,071 genome assemblies (x-axis on a log_10_ scale), (top plot) the regression slope *alpha* (blue line) was estimated as 0.8231 such that the median number of new genes (y-axis) added per isolate (middle plot) was 2.1 (green points) and (bottom plot) the relative rate of new genes (red points) became constant once the number of isolates was >250 (or 10^2.4^). The rate of new genes (green points) and rate of change of new genes (red points) were fitted by loess curves with a span of 0.1 and a degree of two on the average number of new genes per isolate generated by Roary results.

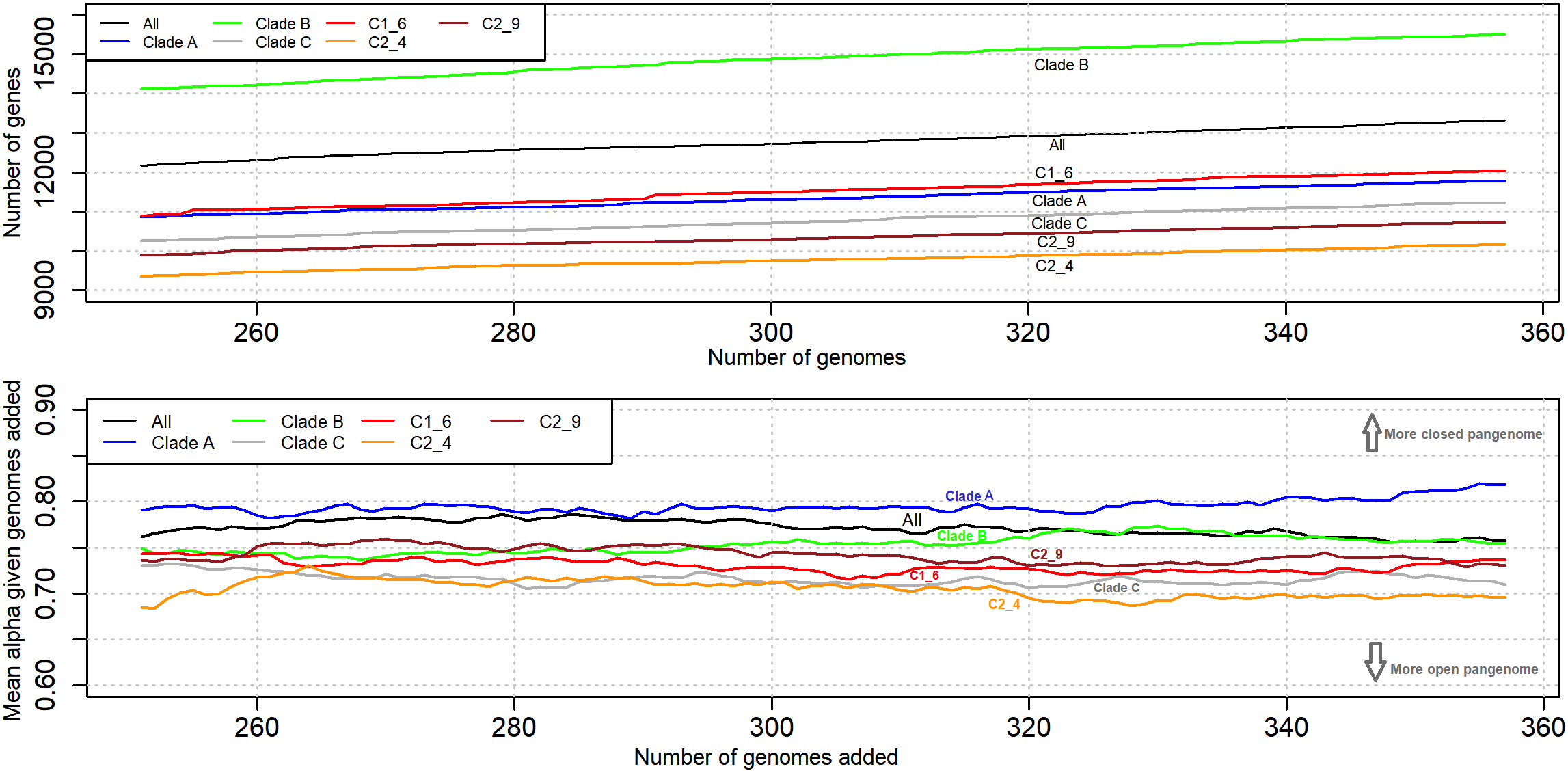

**Supplementary Fig. S12.** The variation in the number of genes and average *alpha* (y-axes) versus the numbers of genomes sampled (shown here for 251 to 357 genomes sampled, excluding missing data) (x-axis). Top: The number of genes added increased gradually across groups with the numbers of genomes, though with higher net diversity of genes in Clade B (green), next C1_6 (red), then Clade A (blue), followed by Clade C (grey), then by C2_9 (brown), and lastly C2_4 (orange). Bottom: The average *alpha* estimated varied with numbers of genomes showed a consistently more open genome for C2_4 (orange, average *alpha* = 0.705), followed by C (grey, average *alpha* = 0.716), then C1_6 (red, average *alpha* = 0.731), before C2_9 (brown, average *alpha* = 0.742), next B (green, average *alpha* = 0.753), the whole collection (black, average *alpha* = 0.771) and A (blue, average *alpha* = 0.795).
